# The genetic architecture of recombination rates is polygenic and differs between the sexes in wild house sparrows (*Passer domesticus*)

**DOI:** 10.1101/2023.01.26.525019

**Authors:** John B. McAuley, Bertrand Servin, Hamish A. Burnett, Cathrine Brekke, Lucy Peters, Ingerid J. Hagen, Alina K. Niskanen, Thor Harald Ringsby, Arild Husby, Henrik Jensen, Susan E. Johnston

## Abstract

Meiotic recombination through chromosomal crossing-over is a fundamental feature of sex and an important driver of genomic diversity. It ensures proper disjunction, allows increased selection responses, and prevents mutation accumulation; however, it is also mutagenic and can break up favourable haplotypes. This cost/benefit dynamic is likely to vary depending on mechanistic and evolutionary contexts, and indeed, recombination rates show huge variation in nature. Identifying the genetic architecture of this variation is key to understanding its causes and consequences. Here, we investigate individual recombination rate variation in wild house sparrows (*Passer domesticus*). We integrate genomic and pedigree data to identify autosomal crossover counts (ACC) and intra-chromosomal allelic shuffling (*r̅*_*intra*_) in 13,056 gametes. Females had 1.37 times higher ACC, and 1.55 times higher *r̅*_*intra*_ than males. ACC and *r̅*_*intra*_ were heritable in females and males (ACC h^2^ = 0.23 and 0.11; *r̅*_*intra*_ h^2^ = 0.12 and 0.14), but cross-sex additive genetic correlations were low (r_A_ = 0.29 and 0.32 for ACC and *r̅*_*intra*_). Conditional bivariate analyses showed that all measures remained heritable after accounting for genetic values in the opposite sex, indicating that sex-specific ACC and *r̅*_*intra*_ can evolve somewhat independently. Genome-wide models showed that ACC and *r̅*_*intra*_ are polygenic and driven by many small-effect loci, many of which are likely to act in *trans* as global recombination modifiers. Our findings show that recombination rates of females and males can have different evolutionary potential in wild birds, providing a compelling mechanism for the evolution of sexual dimorphism in recombination.

## Introduction

Meiotic recombination via chromosomal crossing-over is an essential process in sexual reproduction and has a key role in generating diversity in eukaryotic genomes (Coop and Przeworski 2007). Crossing-over can be beneficial: it ensures proper segregation of chromosomes (Koehler et al. 1996), prevents the accumulation of deleterious alleles, and increases the speed at which populations can respond to selection (Hill and Robertson 1966; Felsenstein 1974; Otto and Lenormand 2002). On the other hand, recombination can increase the risk of mutations at DNA double-strand break sites (Halldorsson et al. 2019; Hinch et al. 2023), and can break down favourable allele combinations previously built up by selection (Charlesworth and Barton 1996). Despite these trade-offs, there is extensive variation in recombination rate both within and between chromosomes, individuals, populations, sexes, and species (Myers et al. 2005; Coop and Przeworski 2007; Ritz et al. 2017; Stapley et al. 2017). The cost-benefit dynamic of recombination rate is likely to vary depending on mechanistic and evolutionary contexts, and if rates are heritable (i.e., there is underlying additive genetic variation), then they have the potential to respond to selection. Therefore, understanding the genetic basis of recombination rate variation - that is, the amount of additive genetic variation and the effect sizes and distributions of gene variants that contribute to such variation - is a critical first step in determining if and how they are contributing to adaptation, and how they themselves are evolving (Ritz et al. 2017; Stapley et al. 2017).

To date, studies investigating the genetic basis of recombination rate variation have mostly been limited to model systems and livestock and have demonstrated that crossover rates can be heritable and associated with particular genetic variants (Cirulli et al. 2007; Smukowski and Noor 2011; Cattani et al. 2012; Chan et al. 2012; Kong et al. 2014; Hunter et al. 2016; Johnston et al. 2016; Kadri et al. 2016; Petit et al. 2017; Johnston et al. 2018; Weng et al. 2019; Samuk et al. 2020; Johnsson et al. 2021; Brekke, Johnston, et al. 2022; Brekke et al. 2023). In some systems such as mammals, the genetic architecture can be oligogenic, with loci with meiotic functions being repeatedly implicated (e.g. *RNF212*, *RNF212B*, *MEI1*, *MSH4, PRDM9*); whereas in other systems, such as Atlantic salmon, variation can be polygenic (i.e., controlled by many loci of small effect). Nevertheless, the patterns observed in these studies may not be generalisable to other systems due to a number of specific factors, such as: relatively low individual variation in recombination rates (e.g. in *Drosophila spp.*); long-periods of female meiotic arrest (e.g. in humans; MacLennan et al. 2015); dynamic variation in karyotype that can affect crossover distributions (e.g. in mice; Capilla et al. 2014; Garagna et al. 2014); and a history of strong and sustained directional selection, which theoretically imposes indirect selection for increased recombination in small populations (e.g. in crops and livestock; Otto and Barton 2001; Ross-Ibarra 2004; but see Muñoz-Fuentes et al. 2015). Another consideration is that in many vertebrates, the positioning of recombination hotspots is determined by *PRDM9*, a gene coding for a zinc finger protein that binds to particular sequence motifs in the genome to initiate double strand breaks (Baudat et al. 2010; Baker et al. 2017). As double strand break repair is mutagenic (Hinch et al. 2023), this can erode the recognised motifs and reduce the number of binding sites, which may in turn select for novel *PRDM9* zinc finger alleles and a corresponding rapid turnover of hotspots (Myers et al. 2010; Grey et al. 2018). Indeed, *PRDM9* is one of the fastest evolving genes in species where it is functional, but it has also been lost from clades such as birds, canids and amphibians, where recombination hotspots appear to be stable and enriched at functional elements (Singhal et al. 2015; Baker et al. 2017). Therefore, research in both non-model, non-*PRDM9*, and non-mammalian systems is required to better understand the genetic architecture of recombination rate more broadly, and how it is affected by other intrinsic and extrinsic variation in natural environments.

A near-universal feature of recombination is that rates and landscapes differ in their degree and magnitude between male and female gametes, a phenomenon known as “heterochiasmy” (Lenormand and Dutheil 2005; Sardell and Kirkpatrick 2020). Numerous hypotheses have been proposed to explain how it has evolved, including: reducing or increasing crossing-over in the sex under stronger selection (Trivers 1988; Burt et al. 1991); gametic selection reducing crossovers in the sex with stronger haploid selection (Nei 1969; Lenormand 2003; Lenormand and Dutheil 2005); counteracting the effects of meiotic drive (Brandvain and Coop 2012); or that it has arisen through drift (Burt et al. 1991). Yet despite several comparative investigations of overall rate (Lenormand and Dutheil 2005; Mank 2009; Cooney et al. 2021), there remains little empirical support for these hypotheses. Interestingly, some of the empirical studies above show that the genetic architecture of recombination rate can differ between the sexes, in terms of the large effect loci underpinning them (Kong et al. 2014; Johnston et al. 2016; Johnston et al. 2018; Brekke, Berg, et al. 2022; Brekke, Johnston, et al. 2022), and/or their cross-sex additive genetic correlations being significantly less than one (Johnston et al. 2016; Brekke et al. 2023). Therefore, studies that specifically dissect within and between-sex genetic architectures in diverse species are imperative to gain a full understanding on the molecular mechanisms and evolutionary drivers/constraints contributing to both recombination rate variation and heterochiasmy from the chromosomal to species levels.

Avian systems present a unique opportunity for further developing our understanding of the genetic basis of recombination rate variation and heterochiasmy in the wild. Linkage mapping and cytogenetic studies in birds have shown that despite relatively conserved karyotypes, there is substantial variation in recombination rates, broad-scale landscapes, and heterochiasmy (Figure 1; see Table S1 for a full list of references). In addition, birds lack *PRDM9* and have highly conserved, stable recombination hotspots that are enriched at transcription start sites (TSS) and gene promoter regions, likely due to increased chromatin accessibility (Pan et al. 2011; Singhal et al. 2015; Kawakami et al. 2017; Bascón-Cardozo et al. 2022). A number of wild bird populations have been subject to long-term, individual based studies of their ecology and evolution. As genomic and pedigree data for these populations proliferates, this presents an opportunity to investigate recombination in wild, non-model systems, and ultimately, the relationships between recombination rates, their genetic architectures, and individual fitness components such as reproduction and offspring viability (Johnston et al. 2022).

**Figure 1:**
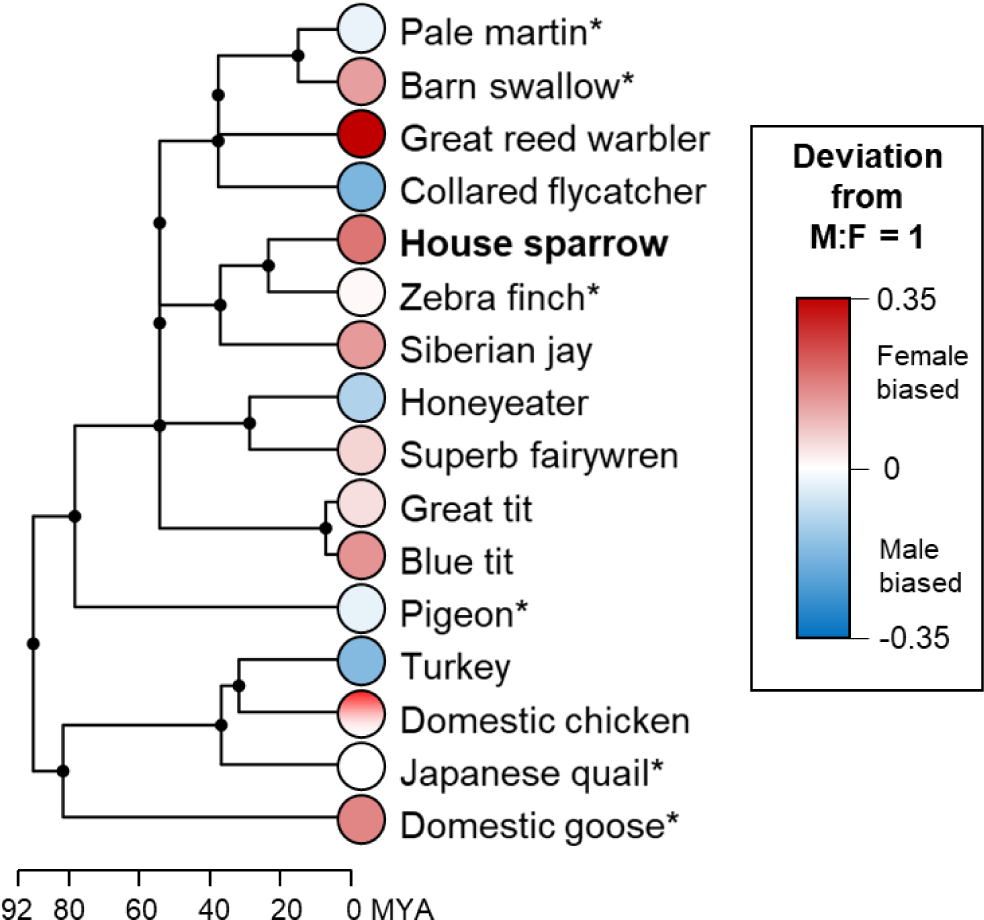
Phylogeny of birds where within-sex recombination rates have been estimated using linkage mapping or cytogenetic analysis of chiasma counts. Chiasma count estimates are indicated with asterisks (*). The underlying data is provided in Table S1 using data compiled in Malinovskaya, Tishakova, et al. (2020) and other sources. The phylogeny was obtained from TimeTree v5 (Kumar et al. 2022). This figure should be taken as illustrative only - some of the linkage map data is likely to have had poor marker densities in telomeric regions and have less resolution to detect double crossovers. As recombination landscape and crossover rates are sex-dependent, this could underestimate recombination in one sex, hence why we have not made a formal comparison. **References:** (Pigozzi and Solari 1998; Pigozzi 2001; Hansson et al. 2005; Calderón and Pigozzi 2006; Backström et al. 2008; Stapley et al. 2008; Groenen et al. 2009; Jaari et al. 2009; Aslam et al. 2010; Hansson et al. 2010; Kawakami et al. 2014; van Oers et al. 2014; Torgasheva and Borodin 2017; Weng et al. 2019; Malinovskaya, Tishakova, et al. 2020; Peñalba et al. 2020; Robledo-Ruiz et al. 2022).

Here, we examine sex-specific variation in individual autosomal recombination rate and its genetic architecture in a wild meta-population of house sparrows (*Passer domesticus*). House sparrows are small passerine birds native to Eurasia, and are now distributed across most human-habited areas of the world. They are human commensals, often found in cities and farmland, and their ubiquity makes them a model system for ecology, evolution and adaptation (Anderson 2006). House sparrows in the Helgeland archipelago have relatively low dispersal from their natal island (Saatoglu et al. 2021), allowing for detailed data on survival and reproduction to be collected since 1993 (Jensen et al. 2004; Stubberud et al. 2017; Araya-Ajoy et al. 2021). This population has extensive genomic resources, including an annotated reference genome (Elgvin et al. 2017), a 6.5K SNP linkage map (Hagen et al. 2020), high-density SNP arrays (Lundregan et al. 2018) and a dense genetic pedigree (Niskanen et al. 2020). In this study, we characterise individual recombination rates using SNP and pedigree data, focussing on autosomes to allow a direct comparison between recombination in the same genome within each sex. We investigate: (a) how recombination varies at the broad scale across chromosomes; (b) the additive genetic variance and genetic architecture of individual recombination rates; and (c) how these factors vary between the sexes, including the potential for female and male rates to evolve independently.

## Results

### Linkage mapping

Autosomal linkage maps were constructed using data from 1,320 full-sib families, incorporating linkage information from 9,777 gametes transmitted by 1,805 individuals to their offspring. We mapped 56,765 autosomal SNPs of known position relative to the house sparrow genome Passer_domesticus-1.0 to 45,022 unique centiMorgan (cM) positions on 28 autosomes, with heterogeneity in the landscape of recombination within and between the sexes (Figures S1 and S2, Tables S2 and S3). The female and male autosomal maps were 1994.4 cM (2.18 cM/Mb) and 1627.2 cM (1.78 cM/Mb), respectively. The female map length was 1.23 times longer than the male map length. There was a strong correlation between chromosome length and genetic map length in each sex (*r*^2^ = 0.97 and 0.96 for females and males, respectively; P < 0.001) and recombination rates were female-biased for most chromosomes (Figure 2A). Chromosome-wide recombination rate (cM/Mb) was higher on smaller autosomes (*r*^2^ = 0.749, P < 0. 0001, fitted as a logarithmic function), with micro-chromosomes demonstrating recombination rates 3 to 5 times that of the genome-wide average (Figure 2B). Despite their high recombination rates, chromosomes 21, 22, 25, 27 and 28 still had linkage map lengths much shorter than the expectation of 50 cM, indicating that the obligate crossover was not always sampled on those chromosomes with the current genomic dataset. Summary statistics for map lengths are provided in Table S2, and the full sex-specific linkage maps are provided in Table S3.

**Figure 2.**
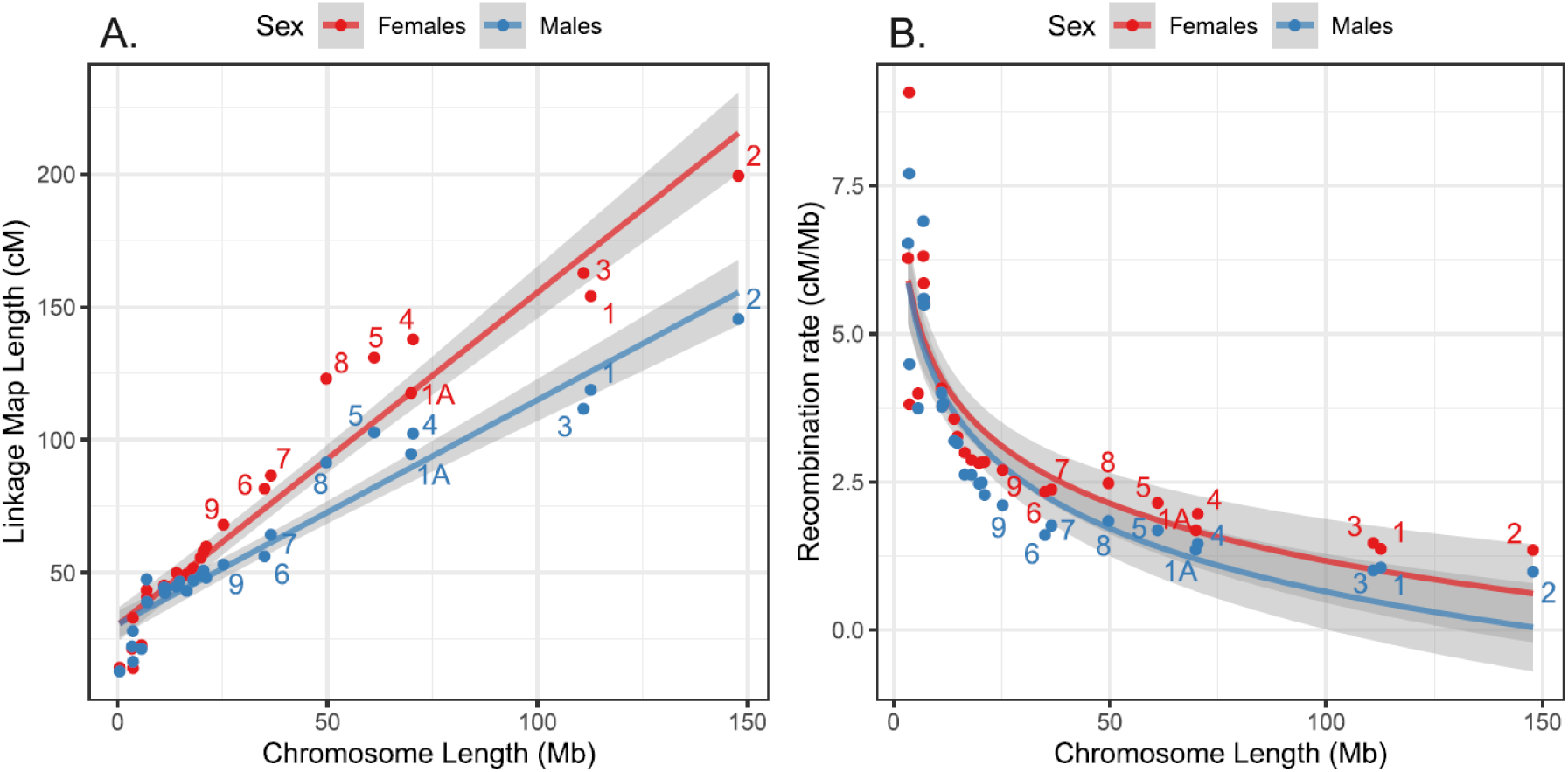
Variation in recombination rates between chromosomes, showing correlations between (A) the sex-specific linkage map lengths (cM) and chromosome length (Mb); and (B) sex-specific chromosomal recombination rate (cM/Mb) and chromosome length. Numbers are individual chromosomes, and lines and the grey-shaded areas indicate the regression slopes and standard errors, respectively. Chromosome 25 has been omitted from Figure 2B due to its exceptionally high recombination rate (30 cM/Mb and 26.9 cM/Mb in females and males, respectively).

### Identification of meiotic crossovers

We used the full pedigree of 12,959 genotyped individuals to phase chromosomes infer crossover (CO) positions in gametes transmitted from focal individuals (FIDs) to their offspring. In total, we identified 236,671 COs in 13,054 phased genotypes from 6,409 gametes transmitted from 1,354 unique female FIDs and 6,647 gametes transmitted from 1,299 unique male FIDs. Larger chromosomes had more COs per gamete than small chromosomes, with chromosomes 1 and 2 having up to 8 COs per gamete in some rare cases in both sexes (Figure 3, Figure S3). Chromosomes 10 to 20 generally had 50% non-recombinant chromosomes, likely reflective of crossover interference over these short chromosomes leading to a single obligate CO, which has a 50% chance of segregation into the gamete. Chromosomes 21 to 28 had more than 50% of chromosomes with no COs, meaning that we had low power to pick up COs on these chromosomes; these chromosomes were discarded from estimates of genome-wide recombination rates.

**Figure 3:**
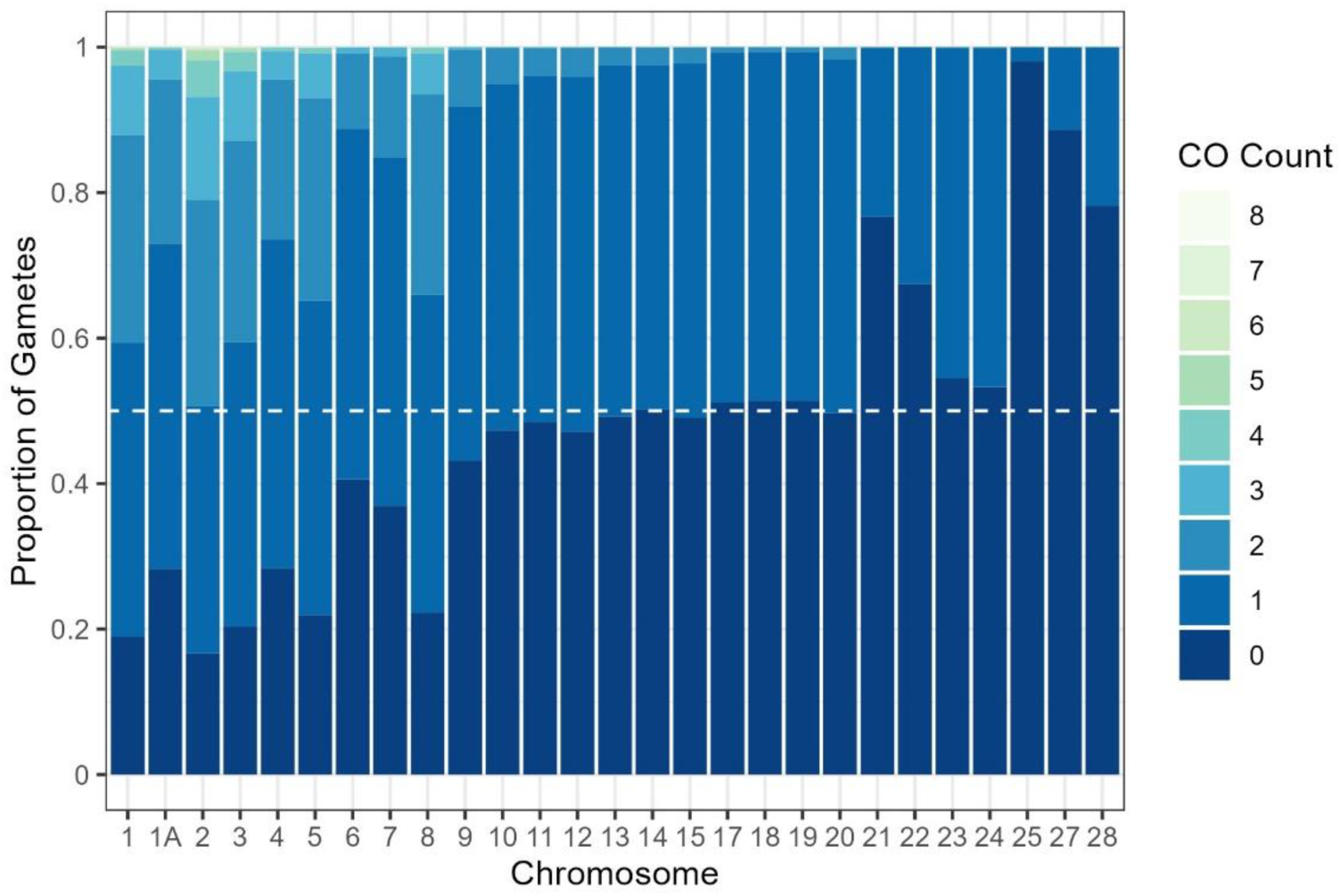
Distribution of crossover (CO) counts per chromosome as the proportion of total number of gametes (N = 13,054). The white dashed line is the minimum expected proportion of gametes with 0 COs per chromosome due to obligate crossing-over and Mendelian segregation of COs into gametes. Separate plots for each sex are provided in Figure S3.

### Individual recombination rate variation

The crossover dataset (for chromosomes 1 to 20 and 1A) was used to calculate recombination rates in each gamete using two measures: the total autosomal crossover count (ACC), and the rate of intra-chromosomal allelic shuffling (*r̅*_*intra*_), which is the probability that two randomly chosen SNP loci on the same chromosome are uncoupled by a crossover during meiosis (Veller et al. 2019). In contrast to ACC, *r̅*_*intra*_ accounts for the effect of crossover positioning, where crossovers located toward the centre of a chromosome will lead to higher shuffling than crossovers located at chromosome ends. The mean ACC was 1.37 times higher in females than in males, with mean ACCs of 18.9 and 13.8, respectively (Figure 4). The mean rate of intra-chromosomal shuffling *r̅*_*intra*_was 1.55 times higher in females (Figure 4). ACC and *r̅*_*intra*_were positively correlated (r^2^ = 0.684, P < 0.0001; Figure S4A).

**Figure 4.**
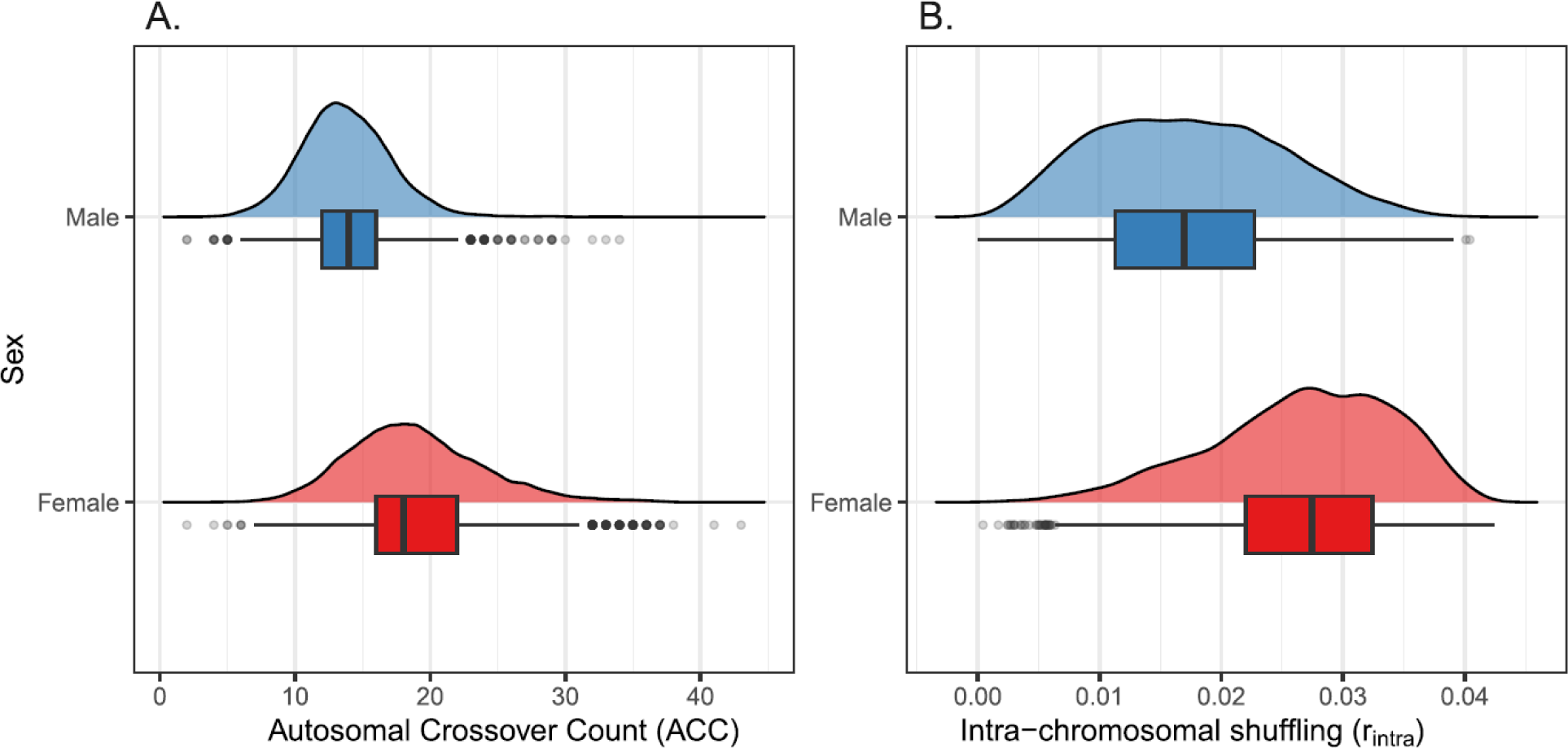
Distributions of female and male recombination rates in gametes transmitted from FIDs to their offspring for A) ACC (autosomal crossover count); and B) *r̅*_*intra*_, (rate of intra-chromosomal shuffling).

### Heritability and genetic correlations of individual recombination rates

The proportion of phenotypic variance in ACC and *r̅*_*intra*_ explained by additive genetic effects (the narrow-sense heritability, h^2^) was determined within each sex using an animal model approach fitting a genomic relatedness matrix as a random effect. We also calculated the mean-standardised additive genetic variance for ACC, defined as the evolvability (*I_A_*), which quantifies the expected proportional change per one unit of selection (Hansen et al. 2011). For all models of ACC and *r̅*_*intra*_, only fixed effects of total phase coverage, and total phase coverage^2^ were significant (Table S4), as were the additive genetic, permanent environment (repeated measures) and residual random effects (Table 1). ACC was heritable and evolvable in females (h^2^ = 0.232, *I_A_* = 0.016) and males (h^2^ = 0.112, *I_A_* = 0.006, P < 0.001, Table 1). The cross-sex additive genetic correlation was positive (r_A_ = 0.292, se = 0.154), but was not significantly different from zero (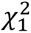 LRT = 3.48, P = 0.062) and was significantly lower than one (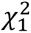 LRT = 19.06, P < 0.001), indicating that the genetic basis of ACC has a significant unshared component between males and females. This was confirmed when conditioning the within sex ACC on the genetic value in the opposite sex, where female and male ACC remained independently heritable (h^2^ = 0.126 and 0.077, respectively, P < 0.001, Table 1). We also calculated the permanent environment effect, a repeated measures parameter which accounts for constant differences between individuals over an above the heritability (Kruuk 2004). This effect was lower but significant in both females and males (pe^2^ = 0.057 and 0.077 respectively, P < 0.05, Table 1), indicating that ACC is also partially explained by individual differences not attributed to additive genetic effects.

**Table 1:** Proportions of phenotypic variance explained by additive genetic (h^2^), permanent environment effect (pe^2^) and residual variance (r^2^). V_P_ and V_P_ (obs) are the phenotypic variances as estimated from the animal model and from the raw data, respectively. The mean value is from the raw data. *IA* is the evolvability (note that *IA* cannot be estimated for ***r_intra_*** due to it being on the absolute scale). h^2^_indep_ is the trait heritability independent of the genetic value of the other sex, and r_A_ is the additive genetic correlation. Values in parentheses are the standard errors. Significances of h^2^, pe^2^ and h^2^_indep_ estimates are indicated by * (P < 0.05), ** (P < 0.001) and *** (P < 0.001). All results are from 6,409 gametes from 1,354 females and 6,647 gametes from 1,299 males on all crossovers occurring on chromosomes 1A and 1 to 20.

| Rate Measure | Sex | $V_P$ | $V_P$ (Obs) | Mean | $V_A$ | $h^2$ | $pe^2$ | $I_A$ | $h^2_{indep}$ | $r_A$ |
| --- | --- | --- | --- | --- | --- | --- | --- | --- | --- | --- |
| ACC | F | 23.93<br>(0.56) | 24.41 | 18.92 | 5.561<br>(0.761) | 0.232<br>(0.029)*** | 0.057<br>(0.023)* | 0.016<br>(2.13e<sup>-3</sup>) | 0.126<br>(0.023)*** | 0.292<br>(0.154) |
|  | M | 10.76<br>(0.221) | 10.56 | 13.76 | 1.202<br>(0.273) | 0.112<br>(0.025)*** | 0.077<br>(0.023)* | 6.35e<sup>-3</sup><br>(1.44e<sup>-3</sup>) | 0.077<br>(0.019)*** |  |
| $\bar{r}_{intra}$ | F | 5.50e<sup>-5</sup><br>(1.04e<sup>-6</sup>) | 5.56e<sup>-5</sup> | 0.027 | 6.40e<sup>-6</sup><br>(7.24e<sup>-7</sup>) | 0.116<br>(0.012)*** | 1.28e<sup>-5</sup><br>(2.43e<sup>-7</sup>) | - | 0.077<br>(0.01)*** | 0.319<br>(0.151) |
|  | M | 5.77e<sup>-5</sup><br>(1.18e<sup>-6</sup>) | 5.75e<sup>-5</sup> | 0.017 | 8.07e<sup>-6</sup><br>(1.43e<sup>-6</sup>) | 0.14<br>(0.027)*** | 0.049<br>(0.020)* | - | 0.082<br>(0.019)*** |  |

*r̅*_*intra*_ was heritable in females and males (h^2^ = 0.116 & 0.140, respectively, P < 0.001, Table 1); as this metric is on the absolute scale, evolvabilities cannot be calculated (see Hansen et al. 2011). When corrected for ACC by adding it as an additional fixed covariate, heritabilities were reduced but remained significant (h^2^ = 0.042 & 0.084 respectively, P < 0.001). The cross-sex additive genetic correlation of ACC was positive (r_A_ = 0.319, se = 0.151) and was significantly different from zero (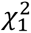 LRT = 4.54, P = 0.033) and one (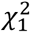 LRT = 23.21, P < 0.001), again indicating that the genetic basis of *r̅*_*intra*_ has an unshared component between males and females. This was confirmed when conditioning the within-sex *r̅*_*intra*_ on the genetic value in the opposite sex, where female and male *r̅*_*intra*_ were independently heritable (h^2^ = 0.082 and 0.077, respectively, P < 0.001, Table 1). There was a significant permanent environment effect for *r̅*_*intra*_ in males (pe^2^ = 0.049), but not females (Table 1).

For both ACC and *r̅*_*intra*_, we investigated the potential contribution of female-restricted chromosomes i.e., the germline-restricted chromosome (Borodin et al. 2022) and the W chromosome to phenotypic variation by fitting individual matriline (i.e., the path of inheritance from mothers to offspring) as an additional random effect. Matriline did not explain any of the phenotypic variance, indicating that neither female-restricted chromosomes significantly contributed to variation in recombination rates in this population. A full explanation of why this analysis was carried out is provided in the Discussion and Methods sections.

### Genome-wide association studies of individual recombination rates

Genome-wide association studies were carried out using a larger genomic dataset of 65,840 SNPs, including Z-linked SNPs and any SNPs of unknown position relative to the house sparrow genome. These models accounted for repeated measures within the same individuals. We did not identify any loci that were significantly associated with variation in ACC or *r̅*_*intra*_ in females or males (Figure 5, Figure S5), with a power analysis indicating that we had 95% power to identify any loci explaining >3.2% of the phenotypic variance. We carried out Empirical Bayes analyses of the GWAS summary statistics to identify loci with significant non-zero effects. For female ACC, 4,696 SNPs (7.13%) and 1,178 SNPs (1.79%) had non-zero effects at the significance thresholds of α = 0.05 and 0.01, respectively (Figure S6). For male ACC, 179 SNPs (0.27%) and 7 SNPs (0.01%) had non-zero effects on ACC at the levels of α = 0.05 and 0.01, respectively (Figure S6). For female *r̅*_*intra*_, 1213 SNPs (1.84%) and 143 SNPs (0.22%) had non-zero effects at the significance thresholds of α = 0.05 and 0.01, respectively (Figure S6). For male *r̅*_*intra*_, 1254 SNPs (1.9%) and 151 SNPs (0.23%) had non-zero effects on ACC at the levels of α = 0.05 and 0.01, respectively (Figure S6).

**Figure 5:**
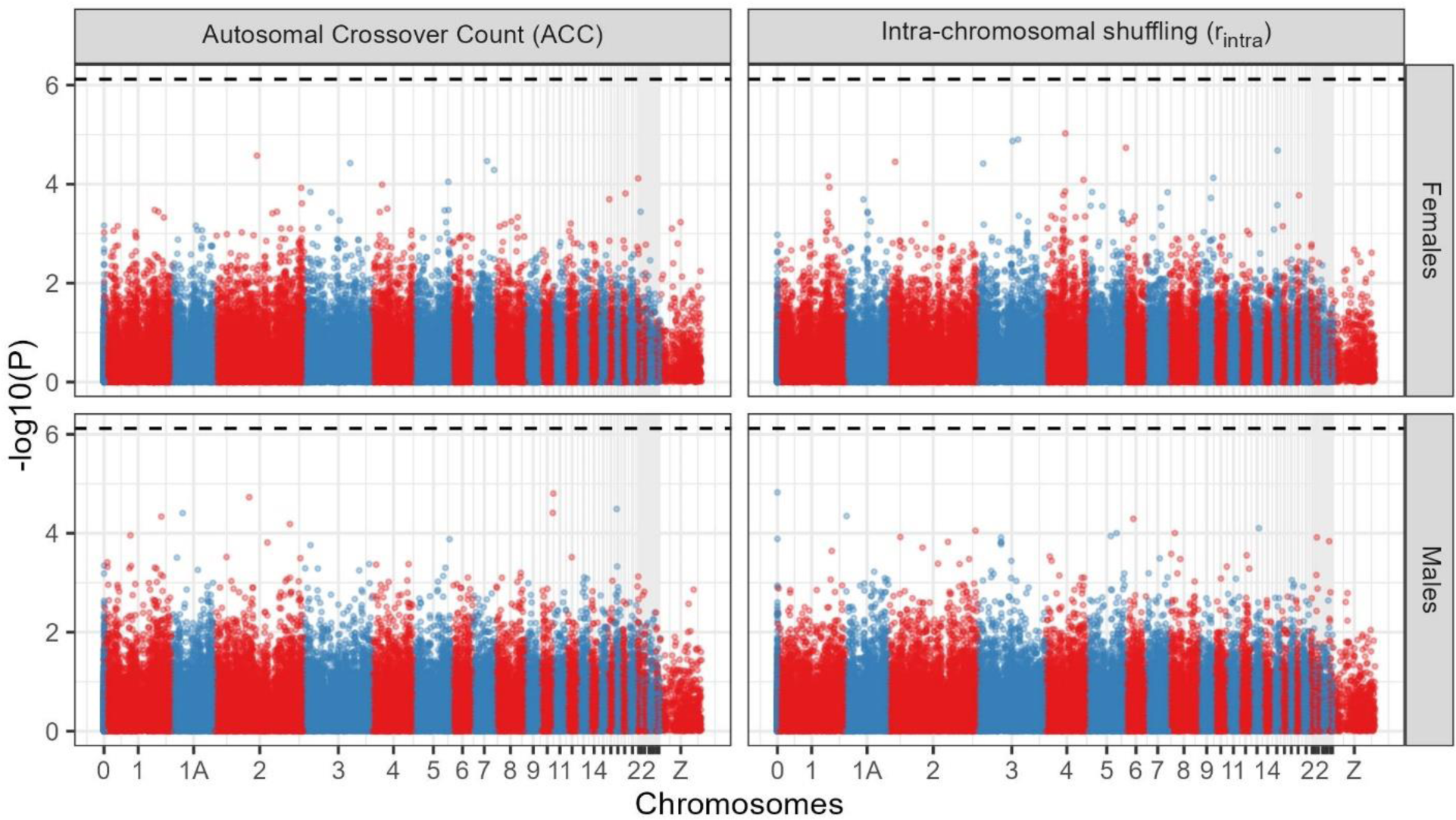
Genome-wide association plots for autosomal crossover count (ACC) and intra-chromosomal shuffling (*r̅*_*intra*_) in females and males at 65,840 SNPs. Phenotypic measures are from 6,409 gametes from 1,354 females and 6,647 gametes from 1,299 males. The dashed line is the genome-wide Bonferroni-corrected significance level (equivalent to α = 0.05). Chromosome 0 indicates SNPs of unknown genomic position. Association statistics have been corrected with the genomic control parameter λ. P-P plots of the null expectations of each plot is provided in Figure S5. Note that visually, P values on the Z chromosome appear to be lower than on other chromosomes. This is an artefact of over-plotting of much higher marker densities on the other chromosomes, and association statistics are not significantly lower on the Z chromosome.

### Chromosome partitioning of additive genetic variance

To confirm the hypothesis of a polygenic architecture of recombination rates, we used a chromosome partitioning approach estimating the contribution of each chromosome to the phenotypic variance (Yang, Manolio, et al. 2011). We computed separate genomic relatedness matrices for (a) chromosome *i* and (b) all chromosomes excluding *i*, and fit both as random effects in an animal model. Larger chromosomes contributed more to the total additive genetic variance for female ACC, female *r̅*_*intra*_, and male *r̅*_*intra*_ (linear regression P < 0.01, Figure 6, Table S6, Table S7), supporting the hypothesis of a polygenic architecture underpinning these traits. The absence of an effect for male ACC is likely due to a lack of model convergence leading to zero estimates for two of the largest chromosomes, 2 and 3; when removing these two chromosomes, the linear regression was significant (t = 4.120, P < 0.001, Adjusted R^2^ = 0.5).

**Figure 6:**
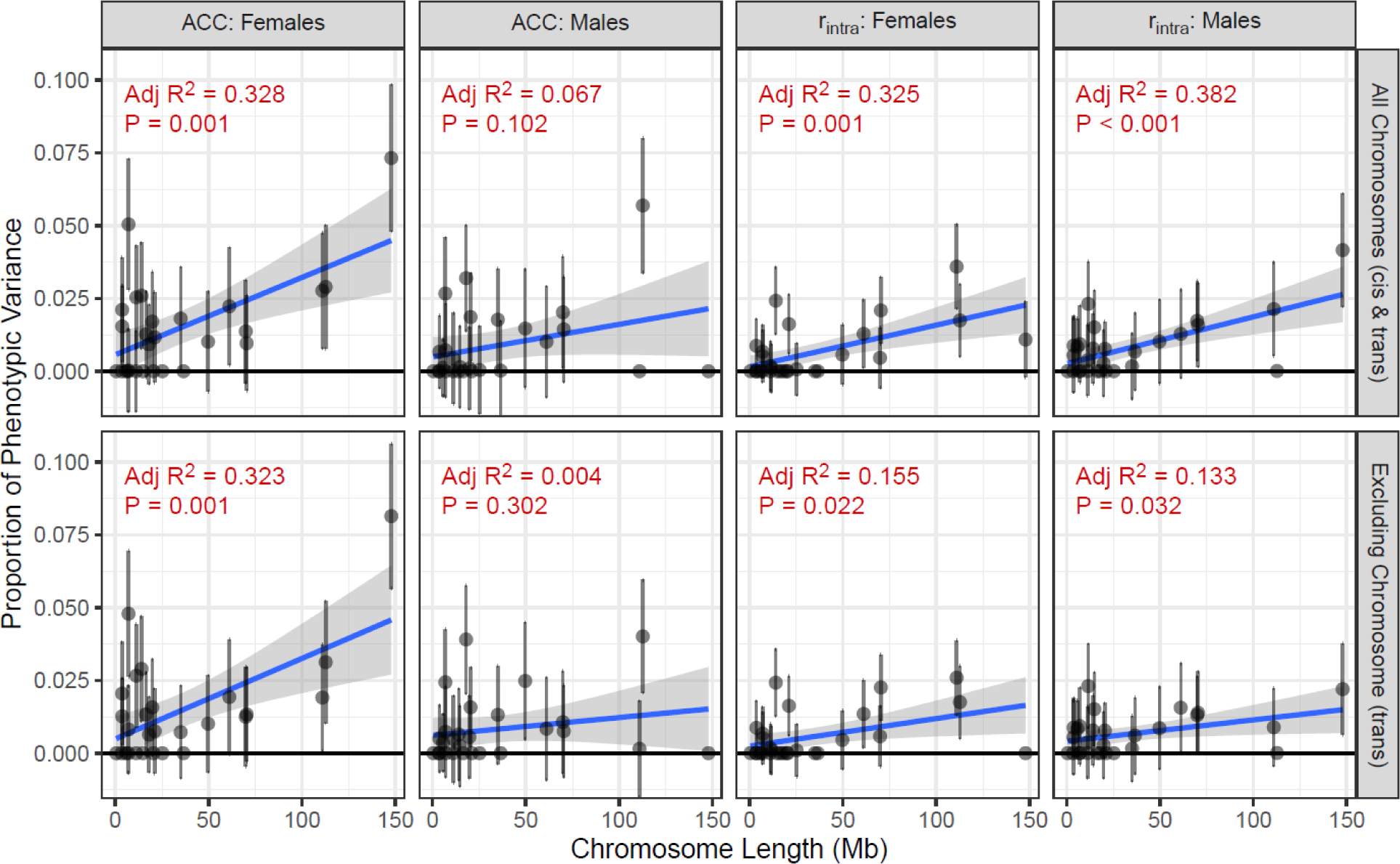
Chromosome partitioning of additive genetic effects on autosomal crossover count (ACC) and intra-chromosomal shuffling (*r̅*_*intra*_) in females and males. Each point indicates the proportion of phenotypic variance explained by a chromosome-specific genomic relatedness matrix modelled as a function of chromosome length. The top row partitions variance for recombination rates estimated on all chromosomes (1-20 & 1A), which accounts for *cis* and *trans* effects combined. The bottom row partitions variance for recombination rates excluding that chromosome, accounting for *trans* effects only. Correlations are analysed with a linear regression (full results in Table S6). Full chromosome partitioning results are provided in Table S7.

We then modified this approach to determine if the polygenic components underpinning recombination rate are commonly acting in *trans* (i.e., they affect the global recombination rate) rather than *cis* (i.e., they affect the recombination rate on the chromosome on which they are situated, and/or are in linkage disequilibrium with heritable aspects of chromatin structure that affect local recombination rates). For each chromosome *i*, we calculated ACC and *r̅*_*intra*_ for chromosomes 1A and 1 to 20 excluding *i*, and then reran the animal models and correlations as above. Similarly, larger chromosomes contributed more to the total additive genetic variance for female ACC, female *r̅*_*intra*_, and male *r̅*_*intra*_ (linear regression P < 0.05, Figure 6, Table S6, Table S7). These effects were weaker for female *r̅*_*intra*_ and male *r̅*_*intra*_ when compared to the full model above, suggesting that a significant component of variation in intra-chromosomal shuffling may still act in *cis*. However, for female ACC, this effect was highly similar in both models, indicating that polygenic variation in female autosomal crossover count is likely to be more commonly acting in *trans* within this population.

## DISCUSSION

In this study, we have shown that female house sparrows have 1.37 times higher autosomal crossover counts (ACC) and 1.55 times more intra-chromosomal shuffling (*r̅*_*intra*_) than males. ACC was moderately heritable in both females and males (h^2^ = 0.23 and 0.11, respectively), as was *r̅*_*intra*_ (h^2^ = 0.12 and 0.14, respectively). A genome-wide association study found no regions of the genome with a significant effect on recombination rate, with chromosome partitioning and Empirical Bayes approaches supporting a polygenic genetic architecture. Larger chromosomes contributed more additive genetic variation to ACC and *r̅*_*intra*_ in the rest of the genome, with further analysis implying that polygenic effects in female ACC are mostly acting in *trans*, i.e., affecting the global recombination rate. For both ACC and *r̅*_*intra*_, the inter-sex additive genetic correlation was low (r_A_ ∼0.3), and both traits remained heritable within each sex after controlling for the genetic values of the opposite sex, indicating that the genetic architecture can evolve independently within each sex. Here, we discuss in more detail the mechanisms by which polygenic architecture is likely to contribute to variation in recombination rate, how this differs between the sexes, and how the broad scale recombination landscape compares with other studies.

### Sex differences in recombination and its genetic architecture

This study provides a compelling example of heterochiasmy in both recombination rates and the broad-scale recombination landscapes. More importantly, our study shows that these sex differences are also underpinned by different genetic architectures. Male and female recombination rates are likely to have some degree of shared genetic architecture, as shown by additive genetic correlations of r_A_ ≈ 0.3; whilst there were moderate standard errors around these estimates, this value was significantly lower than 1 for both ACC and *r̅*_*intra*_. Both traits remained heritable within each sex, even after conditioning on genetic values in the opposite sex, with values ranging from h^2^ = 0.077 to 0.13.

This indicates that there is potential for evolution of recombination rates within each sex independent of that in the other sex, which could affect the degree and potentially the direction of heterochiasmy within the population. With the exception of domestic chickens (Weng et al. 2019), our study is the largest study of individual variation in recombination in birds, and our study is the largest to investigate sex-specific effects at the individual and population levels. As the number of gametes from males and females are similar in our study, and coverage is high across the chromosomes we analysed here, we are confident that we capture the vast majority of biologically meaningful autosomal COs within each sex. While we cannot directly relate heterochiasmy in a single population to a particular hypothesis outlined in the introduction, our results add to a growing body of data that there is sexual dimorphism in not only the rate of crossing over, but also in its positioning and in its genetic architecture. More broadly, our results suggest that while there are fundamental similarities in recombination between female and male meiosis, the observed differences in our study may indicate some distinct biological processes at work between the sexes. Future studies investigating the evolutionary causes and consequences of recombination should endeavour to consider how sex-differences in meiotic processes contribute to observed variation.

### Variation in crossover count and intra-chromosomal shuffling

Much of the theory proposed to explain the advantages and disadvantages of recombination rate variation centres around the generation and preservation of beneficial haplotypes through shuffling of alleles at linked sites (Hill and Robertson 1966; Felsenstein 1974; Kondrashov 1988). The efficacy and extent to which crossovers shuffle linked alleles within chromosomes is not only a function of CO count, but also of the CO position (Veller et al. 2019). For example, a crossover in close proximity to a chromosome end will lead to much less allele shuffling than one in the centre of a chromosome (Veller et al. 2019). However, almost all previous studies investigating the genetic architecture of recombination rates have focussed solely on autosomal crossover counts (e.g. Kong et al. 2014; Ma et al. 2015; Johnston et al. 2016; Petit et al. 2017; but see Brekke et al. 2023). In our study, we modelled both ACC and *r̅*_*intra*_ to consider two different (but not necessarily independent) phenomena, that respectively represent more the variation in the mechanistic process of CO formation (ACC) and the variation in crossover positioning and potentially the evolutionary consequences of recombination (*r̅*_*intra*_). We identified a substantial correlation between ACC and *r̅*_*intra*_ (r^2^ = 0.684), reflecting that much of the variation in *r̅*_*intra*_ is driven by ACC, whereas the remainder may be due to other factors, such as individual differences in chromatin landscape, differences in the genetic architecture, and/or random processes that affect CO positioning. Females demonstrate higher ACC and *r̅*_*intra*_ than males, but this difference is stronger in terms of allele shuffling rather than the number of COs (∼1.5× vs 1.37×, respectively). Therefore, females may exhibit a stronger capacity to drive responses to selection than males in this population, although in reality, it may be that the sex-averaged rate is more meaningful in terms of longer term responses to selection (Burt et al. 1991).

One limitation of our method to characterise COs using pedigree information is that we only use data from gametes that resulted in an offspring. This leads to a “missing fraction” of recombination measures in the population, meaning that our measured rates may not reflect the true rate of crossing over during meiosis; for example, problems in meiosis and lower rates of crossing over can translate into lower fertility and increased rates of aneuploidy in humans (Hassold and Hunt 2001; Kong et al. 2004; Handel and Schimenti 2010; MacLennan et al. 2015). A full understanding of any missing fraction would aim to characterise variation at the pre-zygotic stage e.g. by verifying rate variation using chiasma count data (Malinovskaya, Tishakova, et al. 2020) or gamete sequencing approaches (Dréau et al. 2019) to compare with our pedigree estimated measures. Nevertheless, these methods also have limitations: cytogenetic in birds requires sacrificing individuals to obtain reproductive tissues, and at present, gamete sequencing approaches would be prohibitively expensive to generate large amounts of data (Dréau et al. 2019). Therefore, our pedigree approach is a powerful and useful method to generate large numbers of sex-specific recombination measures (often from pre-existing data in long-term ecological studies).

### What is the nature of polygenic variation underpinning recombination rates?

In our study, we show that recombination rates are heritable and present compelling evidence that this variation is polygenic and driven by many loci of small effect. Indeed, the estimate of h^2^ = 0.232 for female ACC is relatively high compared to other studies of recombination in other vertebrate species (Kong et al. 2014; Johnston et al. 2016; Kadri et al. 2016; Petit et al. 2017; Johnston et al. 2018; Weng et al. 2019; Johnsson et al. 2021; Brekke, Johnston, et al. 2022; Brekke et al. 2023). Our results are similar to mammal studies in that females had higher phenotypic variance and heritability of autosomal crossover counts, but are in contrast in that these studies often identify a conserved suite of loci (e.g. *RNF212*, *RNF212B*, *REC8*, *MSH4*, *HEI10*, among others) that explain a moderate to large effect of heritable variation (e.g. Kong et al. 2014; Kadri et al. 2016; Petit et al. 2017; Johnston et al. 2018; Brekke, Berg, et al. 2022). A lack of significant GWAS in wild populations is often attributed to reduced power to detect trait loci, due to relatively small sample sizes compared to human and livestock studies (Santure and Garant 2018; Johnston et al. 2022). Our power analysis indicated that a locus would have to contribute >3.2% of the phenotypic variance to be detected, meaning that we would only detect moderate to large effect loci. However, we were able to leverage chromosome partitioning and Empirical Bayes approaches to show strong evidence that the additive genetic variance in recombination rates can be attributed to small effect loci throughout the genome, indicating a polygenic architecture. It remains an open question as to how a polygenic architecture of recombination has influenced its evolution. Nevertheless, the maintenance of polygenic variation can arise due to a number of factors, such as a large mutational target for the introduction of new variants (Rowe and Houle 1996), the distribution of selection coefficients over many loci (Sella and Barton 2019), and/or genomic conflict between linked alleles or unmeasured traits with a similar genetic architecture (Teplitsky et al. 2009; Ruzicka et al. 2019).

The question remains - how does polygenic variation contribute variation in recombination rate? This trait ultimately is a phenotype of the genome, and polygenic effects will likely operate through two main routes. First, SNPs may be in LD with genomic features that contribute to local regulation of recombination rate variation in *cis*, e.g. via polymorphic hotspots and heritable aspects of chromatin accessibility that are linked to nearby SNPs. Second, SNPs may be in LD with regions of the genome that affect crossover formation in *trans*, e.g. through modifying the cell environment, chromatin structure, and/or the expression and structure of meiotic proteins. It is difficult to directly test *cis* effects in isolation; this could feasibly test if local variation is associated with local recombination rate. However, as crossovers are inherently rare at a localised level, and with a large number of tests to be conducted, this analysis becomes challenging and underpowered. Nevertheless, we were able to test *trans* effects at a global level by adapting the genome partitioning approach to account for recombination rates on chromosomes excluding the focal chromosome. This indicated that male and female *r̅*_*intra*_ are likely to be affected by both *cis-* and *trans-*acting polygenic variation, whereas female ACC is largely driven by *trans*-acting polygenic variation. Future work will investigate the correlation of non-zero effect SNPs with functional variation, as well as individual and population fine-scale variation of recombination rate to identify common genomic features within the population.

Our study has also shown that the female-restricted chromosomes, namely the W and germline-restricted chromosome (GRC), are likely to make a negligible *trans*-acting contribution to heritable variation in recombination rates within each sex. The GRC is present in all passerines and can comprise around 10% of the genome, but is not present in somatic cells (Pigozzi and Solari 1998; Warren et al. 2010; Torgasheva et al. 2019; Borodin et al. 2022). In males, it is generally ejected before meiosis, whereas in females it duplicates and forms crossovers with itself (Malinovskaya, Zadesenets, et al. 2020; Pei et al. 2022). The function and evolution of the GRC is still poorly understood, but recent work in blue tits (*Cyanistes caeruleus*) has shown that it is enriched for meiotic genes associated with the synaptonemal complex, although its gene content may differ between species (Kinsella et al. 2019; Mueller et al. 2023). Whilst genetic variation on these chromosomes could not be captured by the SNP array (as this only targets somatic genomes), the dense sparrow pedigree allowed us to solve this question by fitting matriline as a random effect. Whilst a negligible effect is likely true in our study population, we cannot rule out that the W or GRC contributes to variation in recombination rate in other systems.

### Trends and comparisons in linkage mapping and broad-scale recombination landscapes

Our motivation for constructing a linkage map was to ensure that the SNPs were correctly positioned relative to one another, as any wrongly positioned SNPs could lead to false calling of crossovers. Our higher-density linkage maps showed strong concordance with a previous 6.5K map in the same population (Hagen et al. 2020). There is an expectation that our higher-density map will have improved resolution at telomeric regions and micro-chromosomes to pick up more COs where recombination rates tend to be higher, and more power to detect double crossovers, leading to longer linkage maps. Contrary to our expectation, the male and female maps were shorter in the current study (∼90% of the previous estimates; Table S1). This is likely due to the use of Lep-MAP3 in the current study, which is less sensitive to phasing or genotyping errors through design (Rastas 2017) compared to the Cri-MAP software (Green et al. 1990) used in the previous study, where such errors can lead to an inflation of the linkage map length. Both studies also use different mapping functions (Morgan vs Kosambi in the current and previous studies, respectively); however, differences in map distances between these functions are negligible in high-density maps, and as marker densities increase, recombination frequencies can be underestimated relative to map distance, which may also explain the observed patterns (Kivikoski et al. 2023).

Our map confirmed heterogeneity in the recombination landscape, particularly between the sexes, where some regions showed strong divergence in rates (e.g. at ∼45Mb on chromosome 1A, or 15-17.5Mb on chromosome 10, among others; Figures S1 and S2). These sex differences indicate that sudden changes in landscape are not necessarily indicative of rearrangements (i.e., when the other sex does not show the same trend), but they could indicate the positions of broader landscape features where males and females can have pronounced heterochiasmy (i.e., differences in male and female rates). Future work will investigate how recombination rate variation, particularly between the sexes, is associated with genomic features. Our overall sex-averaged recombination rate of ∼1.98 cM/Mb was concordant with recombination rates estimated based on cytogenetic analysis of chiasma counts and recombination nodules in birds, which range from 1.6 cM/Mb in female common terns (*Sterna hirundo*, (Lisachov et al. 2017) to 2.9 cM/Mb in female domestic geese (*Anser anser*; Torgasheva and Borodin 2017; see data compiled in Malinovskaya et al. 2018 and Table S1). There is less concordance with previous linkage maps in other species, particularly with those carried out with low marker densities in the early days of linkage mapping, where cM map lengths could be as low as 694 cM for e.g. male Siberian jays (Jaari et al. 2009). It is likely that marker densities in these cases have led to underestimation of recombination by having low sub-telomeric coverage of markers, low sample sizes, and/or markers being too widely spaced to quantify double crossovers. Therefore, understanding how landscapes of recombination vary relative to other avian species requires generation and standardisation of higher density linkage maps in a wider range of systems (Kivikoski et al. 2023).

## CONCLUSIONS

In this study, we have revealed sex differences in the rate and the genetic architecture of recombination in wild house sparrows. This study is an important step in investigating the selective and evolutionary importance of recombination in wild birds, with future analyses focussing on the effects of crossover interference and association between recombination rates and fitness effects at the individual level. Future work will also benefit from a more nuanced understanding of fine-scale variation in recombination from population-scaled estimates, gamete sequencing, and other molecular approaches (Johnston 2024). Integrating this information with our current dataset may shed light on the potential molecular mechanisms underpinning the distribution of recombination (including both crossover and gene-conversion events) and the specific genomic features associated with local variation in heterochiasmy. Overall, our results expand our understanding of individual variation in recombination in a non-mammal system, and our approach has the potential to be extended to other long-term studies with genomic, pedigree, and fitness information.

## Material and Methods

### Study system

All data was collected from the meta-population of house sparrows inhabiting an 18-island archipelago covering 1600 km^2^ off the Helgeland coast in Northern Norway (midpoint of 66°32’N, 12°32’E), which has been subject to an individual-based long-term study since 1993 (Jensen et al. 2004). Birds are routinely captured and individually marked from the beginning of May to the middle of August and for approximately one month in the autumn using mist nets (adults and fledged juveniles), or as fledglings in accessible nests during the breeding season. A small (25 µl) blood sample is collected from the brachial vein for DNA from every captured bird. Individual hatch year was determined as either (a) the first year recorded for nestlings or fledged juveniles captured in the summer and autumn, or (b) the year prior to first year recorded for birds first captured as female adults before June 1 or as males before August 1, or (c) a range including the year first recorded and the year prior for birds first captured as adult females after June 1 or adult males after August 1; hatch island is also recorded alongside hatch year (Ranke et al. 2021; Saatoglu et al. 2021). Sampling was conducted in strict accordance with permits from the Norwegian Food Safety Authority and the Ringing Centre at Stavanger Museum, Norway.

### SNP genotyping and Quality Control

All SNPs used in our analysis were taken from two custom house sparrow Axiom SNP arrays (200K and 70K) based on the resequencing of 33 individual house sparrows (Lundregan et al. 2018). SNPs on both arrays are distributed across 29 autosomes in the house sparrow genome (chromosome 16 was excluded as sequences on this chromosome are difficult to assemble due to containing the highly repetitive major histocompatibility complex). All SNPs on the 70K array are present on the 200K array. All SNP positions are given relative to the house sparrow genome assembly Passer_domesticus-1.0 (GenBank Assembly GCA_001700915.1; (Elgvin et al. 2017; Lundregan et al. 2018)). A total of 3,116 recruited adults were successfully genotyped on the 200K array (Niskanen et al. 2020), and an additional 9,079 recruited adults and non-recruited fledglings and juveniles were successfully genotyped on the 70K array. For our analyses, we merged the two datasets using *--bmerge* in PLINK v1.90b7 (Chang et al. 2015), and removed individuals and SNPs with call rates of < 0.1 (using *--mind* and *--geno*) and SNPs with minor allele frequencies (MAF) of < 0.01 in founder individuals (*--maf*). The merged dataset contained 65,840 SNPs in 12,965 individuals.

For all autosomal SNPs used in the estimation of recombination rates below, we conducted a second, stricter round of quality control to minimise the risk of genotypic and/or phasing errors leading to spurious calls of recombination events. Mendelian errors were identified with the *--mendel* function in PLINK, and SNPs with more than 100 Mendelian mismatches were discarded. It is possible that these mismatches have arisen due to DSB repair via biased gene conversion events (Lorenz and Mpaulo 2022); however, as this study only considers crossover events, there is no loss of relevant information by conducting this quality control step. We generated summary statistics of SNP loci using the function *summary.snp.data* in GenABEL v1.8-0 (Aulchenko et al. 2007) in R v3.6.3. To ensure strong concordance between 200K and 70K data, we removed SNPs where there was a difference between the datasets of more than 3 standard deviations between (i) the minor allele frequency, (ii) deviation from Hardy-Weinberg equilibrium, and/or (iii) the proportion of heterozygote individuals. After these steps, we retained 56,767 autosomal SNPs and 617 Z-linked SNPs in 12,959 individuals. The mean and median autosomal inter-marker distances in this final dataset were 16.1 kb and 9.6 kb, respectively, and the mean and median Z-chromosome inter-marker distances were 111.2 kb and 77.9 kb, respectively. We investigated the linkage disequilibrium (LD) exponential decay profile for all autosomal loci occurring within 500 kb windows using the flag *--ld-window-kb 500* in PLINK. LD decayed to r^2^ = 0.01 at a distance of ∼100 kb (Figure S7).

### Genetic pedigree construction

A metapopulation-level pedigree was constructed using a subset of 873 SNP markers in the software Sequoia (Huisman 2017). This subset of SNPs was selected for the pedigree construction by filtering the 70K SNP set using PLINK 1.9 (Chang et al. 2015) using the *--indep* command, specifying a window size of 1000 KB, step size of 10Kb and an r^2^ threshold of 0.01, whilst filtering for MAF > 0.35 and excluding Z-chromosome mapped SNPs. The SNP set was further refined by constructing a preliminary parentage-only pedigree, and removing SNPs with Mendelian error rates > 0.03. After these steps, 873 SNPs remained. For genetic sex assignment, we calculated the *Fhat2* inbreeding coefficients (F_Z_) using the *--ibc* command using 306 high-quality Z-chromosome SNPs. Individuals with F_Z_ > 0.95 were assigned as female and those with F_Z_ < 0.5 assigned as male. Individuals with F_Z_ > 0.5 < 0.95 and were left as “unknown” sex, as were individuals assigned as genetic females that had autosomal F_ROH_ > 0.1, due to inbreeding biasing F_Z_ estimates. We used the R-package Sequoia (Huisman 2017) to construct a pedigree including 11,073 individuals sampled from Helgeland and several other nearby populations. We used the genetic sex for all individuals and assigned known hatch years as the year of first capture for individuals sampled as nestlings or first captured as juveniles, and assumed sparrows first captured as female adults before June 1 or as males before August 1 were hatched the previous year. We set a possible hatch year range of year *t*-1 or *t* for sparrows first captured as adult females after June 1 or adult males after August 1 in year *t*, because by this stage some juveniles are difficult to distinguish from adults. After the initial parentage-only pedigree, a final pedigree was constructed with sib-ship clustering and LLR calculation enabled, with a maximum of eight sib-ship iterations and a genotyping error rate of 0.002 (Niskanen *et al*. 2020).

### Linkage map construction

Autosomal linkage map construction was conducted using Lep-MAP v3 (Rastas 2017). The pedigree was ordered into full-sib families as follows: for each unique male-female mating pairing (hereafter referred to as the focal individuals, or FIDs, in which meiosis took place), we constructed a three-generation family including all genotyped parents and offspring. Whilst an FID can be present in several families (i.e., as an offspring, parent, or when mating with a different individual), this design meant that each meiosis was only counted once. A total of 4,534 full-sib families were constructed, with 1 to 20 offspring per family. We assumed the same marker order as the house sparrow genome above, and treated each chromosome as a separate linkage group. The module *filtering2* was run with the parameter *datatolerance* = 0.01 to filter based on segregation distortion, with all markers passing this step. The module *separatechromosomes2* was run for each linkage group, and markers that were not assigned to the main group (i.e., LOD score < 5) were excluded (N = 2 SNPs removed). Finally, the module *ordermarkers2* was run to calculate the centiMorgan (cM) positions using the Morgan mapping function for each sex separately. Relationships between (a) chromosome length (in megabases) and linkage map length (in cM) and (b) male and female linkage map lengths (in cM) were analysed using linear regressions in R v4.2.2.

### Calculating individual recombination rates

#### Chromosome phasing and crossover estimation

The software YAPP v0.2a0 (https://yapp.readthedocs.io/) (Servin 2021) was used to phase chromosomes and identify crossover (CO) positions in gametes transmitted from FIDs to their offspring. This approach uses the whole pedigree rather than the smaller sub-pedigrees above, and is more robust to missing individuals, allowing us to characterise crossovers in more individual gametes. YAPP identified 14,769 parent-offspring pairs, representing 14,769 gametes in which COs could potentially be inferred. First, the *mendel* command was run with default parameters, removing 143 pairs with higher rates of Mendelian errors using the default parameters. Next, the *phase* command was used to infer the gametic phase of chromosomes. The *phase* analysis proceeds in two stages through the pedigree, ensuring parents are processed before their offspring. In the first stage, Mendelian transmission rules are applied to reconstruct the gametes passed from parent to offspring (e.g., a homozygous individual can only transmit one of the two alleles to its offspring). With this information, the phase of an individual (i.e., the haplotypes received from each parent) is determined by (i) combining all haplotypes transmitted to its offspring using the Weighted Constraints Satisfaction method of Favier et al. (2010) implemented in ToulBar2 and (ii) the haplotypes transmitted by its parents. In the second stage, the partially reconstructed gametes are used to infer segregation indicators with a Hidden Markov Model, as described in Fledel-Alon et al. (2011) and Druet and Georges (2015). These segregation indicators allow for a more precise reconstruction of the gametes transmitted from parent to offspring, which is then used to produce the final phase reconstruction for all individuals. Finally, the *recomb* command was run to identify COs from the segregation indicators. For each chromosome and each meiosis, YAPP outputs the start and stop positions of the informative length of the chromosome (i.e., the total region where phase can be inferred for a particular individual from the pedigree, or “coverage”) and the start and stop positions of each crossover interval as determined by the Hidden Markov Model (Figure S4A); this was included to account for the uneven information for CO detection across parent/offspring pairs (e.g. due to variation in inbreeding). This process was run in three iterations of the *phase* and *recomb* commands to allow conservative quality control and minimise the risks of calling false COs:

##### Iteration 1

After the first iteration, we removed parent-offspring links with > 60 COs per gamete and/or > 9 COs on any chromosomes, and removed parents or offspring with an autosomal heterozygosity of < 0.36, as this threshold was associated in an uptick in estimated CO counts, suggesting a reduced sample quality and/or increased phasing difficulty (N = 14,127 gametes remaining).

##### Iteration 2

After the second iteration, we investigated the genomic locations of double crossovers (DCOs) in close proximity (< 3Mb between adjacent CO mid-points). Chromosome 26 had a high number of close DCOs despite its short length (∼6.9 Mb) indicating that this chromosome either has many structural variants or is poorly assembled, and so all COs on this chromosome were discarded. We then identified genomic regions enriched for close DCOs by splitting the genome into 100 kb bins and tallying the number of CO window mid-points in each bin. Two regions of the genome showed elevated close DCOs at chromosome ends: these were from 0-3 Mb of chromosome 4; and from 68-69.9 Mb of chromosome 1A (Figure S8). We speculate that these regions have large structural variants (e.g. inversions) and/or rearrangements relative to the reference genome that lead to false calling of DCOs in these regions. In this case, we removed all COs overlapping these two regions from all further analyses. We then removed all parent-offspring links with > 45 COs per gamete and/or > 9 COs on any chromosome, and removed parents and offspring with a SNP call rate of < 0.98 and/or a heterozygosity value outside 3 standard deviations of the mean (N = 13,159 gametes remaining).

##### Iteration 3

After running for a third iteration, we re-investigated all DCOs in the data. We assumed that all COs were Class I crossovers and subject to crossover interference (>90% of COs in vertebrates, see Pazhayam et al. 2021). Therefore, we also assumed short DCOs were indicative of non-crossover gene-conversion events, non-interfering Class-II COs, phasing errors, and/or genotyping errors. After visual examination of the distribution of distances between DCOs to determine an appropriate threshold, we identified an increase in DCOs occurring within an interval of 2Mb or less. Whilst the distance at which crossover interference operates in birds is generally unknown, the lack of DCOs on chromosomes less than 10Mb in length (Figures 2 & 3) indicates that the 2Mb threshold is unlikely to incorporate Class I COs. Therefore, we removed all COs that were less than 2Mb apart (from the right-hand boundary of the first CO interval to the left-hand boundary of the second CO interval; Figure S9B). Any gametes with more than 5 short DCOs were removed. In cases of clustered short COs (i.e., 3 or more adjacent COs that are separated by distances of 2 Mb or less), 1 CO was called in the case of odd numbers of COs (i.e., a phase change occurred at either side of the cluster), or 0 COs in the case of even numbers of COs (i.e., there was no phase change on either side of the cluster).

It should be noted that pedigree-based methods to estimate COs can only identify those present in one of the four cells resulting from meiosis. For each CO, there will be two recombinant and two non-recombinant chromatids at that position. Therefore, our CO counts represent a sample of the crossovers that happened in meiosis I. We assume that each CO is sampled with a 50:50 probability, but we cannot rule out that meiosis with two or more COs on the same chromosome may be more likely to be co-inherited on the same chromatid. Therefore, there is an expectation that at least 50% of gametes will have at least one CO per chromosome due to obligate crossing-over, as each CO per meiosis has a 50% chance of being observed in the gamete due to Mendelian segregation. We observed that the micro-chromosomes 21 to 28 had a higher-than-expected number of gametes with no observed COs (>50%; Figure 3), meaning that not all COs can be detected. Therefore, all COs occurring on these chromosomes were discarded from downstream analyses. In total, we identified 212,711 COs in 13,056 phased genotypes from 6,409 gametes from 1,354 unique females and 6,647 gametes from 1,299 unique males.

#### Recombination rate calculation

The CO dataset was used to calculate recombination rates in FIDs using two approaches. First, we determined the autosomal crossover count (ACC) by summing the number of COs per gamete. Second, we calculated the rate of intra-chromosomal allelic shuffling, *r̅*_*intra*_, which is the probability that two randomly chosen SNP loci on the same chromosome are uncoupled in meiosis (Veller et al. 2019). This was defined as:

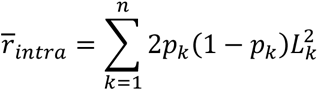

where for chromosome *k*, *p_k_* is the proportion of the SNPs inherited from one parent, *L_k_* is its fraction of the genome length (or gene count), and *n* is the number of autosomes. In our initial data exploration, we also quantified *r̅*_*intra*_ as a function of individual genes rather than SNP loci (defined as *r̅*_*gene*_). However, *r̅*_*intra*_ and *r̅*_*gene*_ were highly correlated (Pearson’s correlation r = 0.948, P < 0.001; Figure S4B) and so only the *r̅*_*intra*_ measure was used in downstream analyses.

### Determining Heritability of Recombination Rate

#### Univariate models

Variance components and the proportion of phenotypic variance in recombination measures attributed to additive genetic effects (the narrow-sense heritability, *h*^2^) were determined using an “animal model” fitted by restricted maximum-likelihood in the package ASReml-R v4 (Butler et al. 2009) in R v4.2.2. A genomic relatedness matrix (GRM) based on all autosomal markers was constructed with GCTA v1.94.1 (Yang, Lee, et al. 2011). The GRM was adjusted for sampling error using the --*grm-adj 0* argument, which assumes the frequency spectra of genotyped and causal loci are similar. Models were run for ACC and *r̅*_*intra*_ in males and females separately. The fixed effect structure included the total phase coverage from YAPP (the length of the genome that can be phased and therefore within which crossovers can be detected) and the total phase coverage squared. For models of *r̅*_*intra*_, we fit with and without ACC as an additional continuous fixed effect, as intra-locus shuffling is a function of the crossover count, with both measures highly correlated (Pearson’s correlation r = 0.684, P < 0.001; Figure S6A). Random effects included the additive genetic effect (GRM) and permanent environment effect. The permanent environment effect is a repeated measures parameter which accounts for constant differences between individuals over and above the additive genetic effect, which can be generated by differences in individual environment and condition, long-term effects of critical developmental stages, and dominance and epistatic genetic effects (Kruuk 2004). A failure to account for this effect can lead to upwardly biased estimates of the additive genetic effect (Kruuk and Hadfield 2007). Models were also run with a pedigree-based relatedness matrix calculated using the *ainv* function in ASReml-R, but variance estimates were highly similar to those from the GRM. Initial models were run with age of the FID in year of gamete formation (defined as the difference between offspring hatch year and parent hatch year) as a continuous fixed covariate, and random effects of FID hatch year, FID natal island and offspring hatch year (to investigate cohort effects and parse apart environment effects) and FID’s mother identity (to estimate maternal effects). However, these effects were estimated as bounded at zero and were not significant in any models, and so were discarded from further analyses.

The heritability of each measure (*h^2^*) was determined as the ratio of the additive genetic variance *V_A_* to the total phenotypic variance *V_P_*, defined as the sum of random effect variances and the residual variance as estimated by the animal model, using the equation *h*^2^ = *V*_*A*_/*V*_*P*_. We also calculated the mean-standardised additive genetic variance, defined as the evolvability (*I_A_*) using the equation *I*_*A*_ = *V*_*A*_/(*x̅*^2^), where *x̅* is the trait mean. This measure quantifies the expected proportional change per one unit of selection (Hansen et al. 2011). Standard errors of these estimates were derived using the delta method implemented in the ASReml-R function *vpredict*.

#### Bivariate models

Bivariate models were run to determine the additive genetic covariances and correlations between male and female ACC and *r̅*_*intra*_ with total phase coverage as a continuous fixed effect. The additive genetic correlation, r_A_, was calculated without constraint using the CORGH function (i.e., correlation with heterogenous variances) in ASReml-R v4. This function also allowed us to run additional models where r_A_ was constrained to zero or 0.999 (as a correlation of one cannot be fit by the software). Significant differences between the observed value and constrained models were tested using likelihood ratio tests, calculated as 2 times the difference between the model log-likelihoods, distributed as 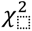 with 1 degree of freedom.

The CORGH function reports the heritabilities of traits when unconditional on the genetic values of the other sex. To confirm this, we also estimated how much of V_A_ in females was conditional on genetic values in males (i.e., *V*_*A*(*f*|*m*)_), using the following equation:

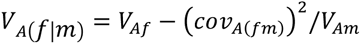

where *V*_*Af*_ and *V*_*Am*_ are the additive genetic variance estimates in females and males, respectively, and *cov*_*A*(*fm*)_ is the additive genetic covariance of the trait between the sexes (Hansen et al. 2003). The conditional heritability was then calculated as 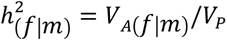. The *cov*_*A*(*fm*)_ estimate was obtained from the same bivariate animal model as above, specifying the US function (i.e., the covariance matrix is unstructured). This was then repeated for males, conditional on genetic values in females (i.e., *V*_*A*(*m*|*f*)_).

#### Potential Contribution of Female-Restricted Chromosomes to Variation

House sparrows have two female-restricted chromosomes present in the cell during meiosis I - a haploid germline-restricted chromosome (GRC; Pigozzi and Solari 1998; Warren et al. 2010; Torgasheva et al. 2019; Malinovskaya, Zadesenets, et al. 2020; Borodin et al. 2022; Pei et al. 2022) and the W chromosome. The function and evolution of the GRC is poorly understood, but recent work in blue tits (*Cyanistes caeruleus*) has shown that it is enriched for meiotic genes associated with the synaptonemal complex, although its gene content may differ between species (Kinsella et al. 2019; Mueller et al. 2023). If there is between-individual genetic variation for recombination rate on the GRC or W chromosomes, this may be partially captured by the GRM (as related females will carry more genetically similar GRCs and W chromosomes), but *not* by the SNP array variation above. To determine their potential contribution to additive genetic variation in recombination, we used the pedigree to determine the matriline identity of each bird (i.e., the path of inheritance from mother to offspring). Mother identity was known for 83% of birds through direct observation or inference of unsampled parents from previous pedigree construction in Sequoia (Huisman 2017; Niskanen et al. 2020). We identified 351 unique matrilines for the 1,354 female birds with recombination rate estimates, with 90% of birds assigned to the 150 most common matrilines. We then fit individual matriline (representing maternal inheritance of GRC and W, as well as mitochondria) as an additional random effect in the animal models above to partition the proportion of variance explained by the line of maternal inheritance independent of the additive genetic effect. If significant, this implies that variation on the GRC, W chromosome, or mitochondria may also contribute to heritable variation in recombination rate measures.

### Determining Genomic Variants Associated with Recombination Rate

#### Genome-wide association studies (GWAS)

GWAS of recombination measures were conducted with all FIDs using the merged SNP dataset (N = 65,840) implemented in RepeatABEL v1.1 (Rönnegård et al. 2016) in R v3.6.3. Models were run for ACC and *r̅*_*intra*_ in males and females separately. The package models the additive effect of SNPs, where genotypes (AA, AB and BB) correspond to 0, 1 and 2, and slopes and standard errors of associations are calculated. The total phase coverage was fit as a fixed effect, and the GRM was fit to account for inflation of test statistics due to population structure. To account for any further inflation, we divided association statistics using the genomic control parameter λ, which was calculated as the observed median χ^2^ statistic, divided by the null expectation median χ^2^ statistic (Devlin and Roeder 1999). The significance threshold was calculated using a Bonferroni correction, with the threshold set at α = 0.05 of P = 2.765 ×10^−7^. We performed a power analysis to evaluate the capacity of our GWAS to detect biologically meaningful quantitative trait loci using the method outlined in (Visscher et al. 2017) implemented in an R function provided by Kaustubh Adhikari (https://github.com/kaustubhad/gwas-power) in R v4.2.2. When specifying a minimum sample size of N = 1299 unique males in our dataset (i.e., the most conservative threshold), we determined that we had 95% power to identify a locus explaining 3.2% of the phenotypic variance.

#### Distribution of polygenic effects

We determined the distribution of allele effect sizes and estimated false discovery rates and false sign rates to identify loci with non-zero effects on recombination rate using the *ash* function in the R package ashR v2.2-32 (Stephens 2017). This package models the slopes and standard errors of the additive SNP effects from the GWAS in an Empirical Bayes framework to compute a posterior distribution of SNP effect sizes across all loci. For SNPs estimated to have non-zero effect on the trait, the significance of a SNP effect is determined by a local false sign rate, defined as probability of error when classifying the slope of the effect as positive or negative, with cut-off thresholds at α = 0.05 and α = 0.01. The prior distribution was specified to be unimodal and symmetric around 0 when applying the false discovery rate estimation, i.e., effect sizes are most likely to be 0 and equally likely to be positive or negative; this was specified using the arguments *mixcompdist = “uniform“* and *method= “fdr“*.

#### Chromosome partitioning of additive genetic variance

We estimated the contribution of each chromosome to the additive genetic variation in recombination rate, to determine if larger chromosomes (i.e., those with more genes) contribute more to the total additive genetic variance, and thus supporting the hypothesis of a polygenic genetic architecture. For each chromosome *i*, we calculated two GRMs: one for chromosome *i* and one for all autosomes excluding *i*. GRMs were determined using GCTA v1.94.1 with the same parameters above. We then fit both GRMs in place of the single GRM in the animal model structure above. This allowed us to determine the proportion of variance of the global recombination rate explained by each chromosome. We then investigated the correlation between chromosome *i* size and the proportion of additive genetic variance explained by chromosome *i* using a linear regression in R v4.3.1. It should be noted that chromosome partitioning analyses can be biased as a result of heteroscedasticity and censoring (Kemppainen and Husby 2018), as the trait heritability, SNP effect sizes, and their physical location will impact inferences of polygenic architecture (Kemppainen and Husby 2018). To quantify the effect of this, we repeated the chromosome partitioning analysis above, but permuting the phenotype values across all individuals within the model. This was done 100 times, due to computational constraints. We compared the permuted data linear regression with the true regression above, identifying little impact of censoring and heteroscedasticity on our current analysis (Figure S10).

Finally, we adapted the chromosome partitioning approach to determine if genetic variants underpinning recombination rate are more likely to commonly act in *trans* (i.e., they affect the global recombination rate) or *cis* (i.e., they affect the recombination rate on the chromosome on which they are situated). More simply, this investigates the contribution of each chromosome to recombination on the remaining chromosomes. We repeated the analysis above, except for each chromosome *i*, we calculated the ACC and *r̅*_*intra*_ response variables excluding measures from chromosome *i*. We then investigated the correlation between chromosome *i* size and the proportion of additive genetic variance explained by chromosome *i* using a linear regression in R v4.3.1, as above.

## DATA STATEMENT

Data will be publicly archived on Dryad for publication. All scripts are available at the following link for review: https://github.com/susjoh/2024_Sparrow_Recomb_GWAS.

## ACKNOWLEDGEMENTS

We thank students and fieldworkers for help with fieldwork, and the hospitality of inhabitants at the study area in Helgeland who made this study possible. We thank Katie Abson, Jarrod Hadfield, Anna Hewett, Mark Ravinet, Martin Stoffel, Alexander Suh, Anna Torgasheva, and Carl Veller for helpful discussions on many aspects of the study. This work made extensive use of the Ashworth Computing Cluster Co-operative (AC3) hosted at the Institute of Ecology and Evolution, with assistance from Dominik Laetsch, Cei Abreu-Goodger, and Jobran Chebib.

## FUNDING

The house sparrow field study and genomic resource development was funded by the Research Council of Norway (RCN grant nos. 221956, 274930 and 302619), the RCN’s Centres of Excellence funding scheme (grant no. 223257) and The Royal Society (RGF/R1/180090). S.E.J. was funded by a Royal Society University Research Fellowship (UF150448 and URF/R/211008). J.B.M. was funded by a Darwin Trust of Edinburgh PhD studentship.

## AUTHOR CONTRIBUTIONS

SEJ & HJ conceived and designed the study. HJ and THR organised and collected field data together with AH, HAB and IJH. HAB, HJ, AH, IJH & AKN generated and curated the genomic dataset. HAB & AKN constructed the pedigree. BS developed and adapted YAPP software for the analysis. JBM, CB, LP & SEJ analysed the data. SEJ & JBM wrote the manuscript with input from all authors.

## Notes

### Competing Interest Statement

The authors have declared no competing interest.

### Summary of Updates

The title of the paper was modified to better reflect our findings. Some edits and clarifications were made to the methods and results of the paper. A new supplementary figure was added demonstrating variation in recombination in the recombination landscapes. No changes were made to in the analyses and all conclusions are the same as the previous revision.

