## Supplementary Figures and Tables for "The genetic architecture of recombination rates is polygenic and differs between the sexes in wild house sparrows (*Passer domesticus*)"

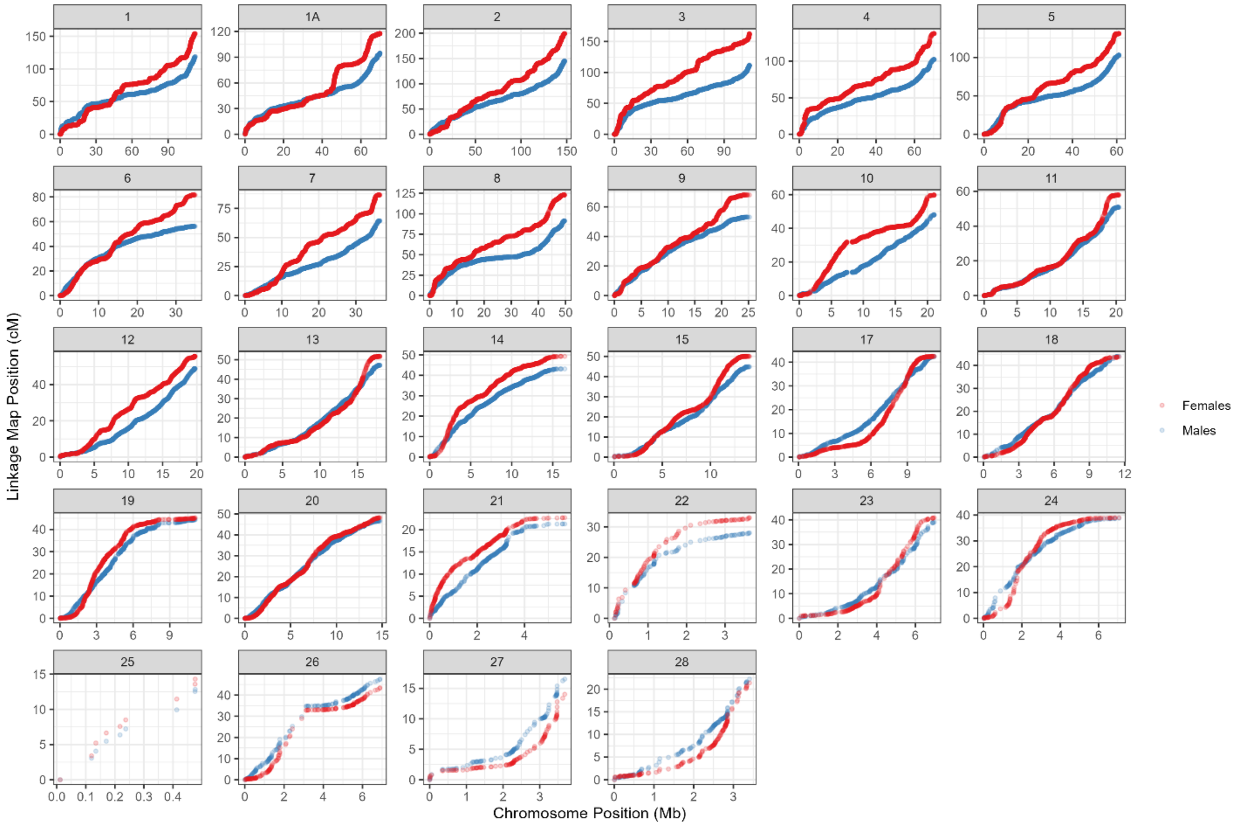


**Figure S1:** Sex-specific autosomal genetic linkage maps for house sparrows, with centiMorgan (cM) position shown relative to the chromosome position (in Mb) on the house sparrow genome assembly Passer_domesticus-1.0 (Elgvin et al. 2017). Map summary data are provided in Tables S2 and S3.

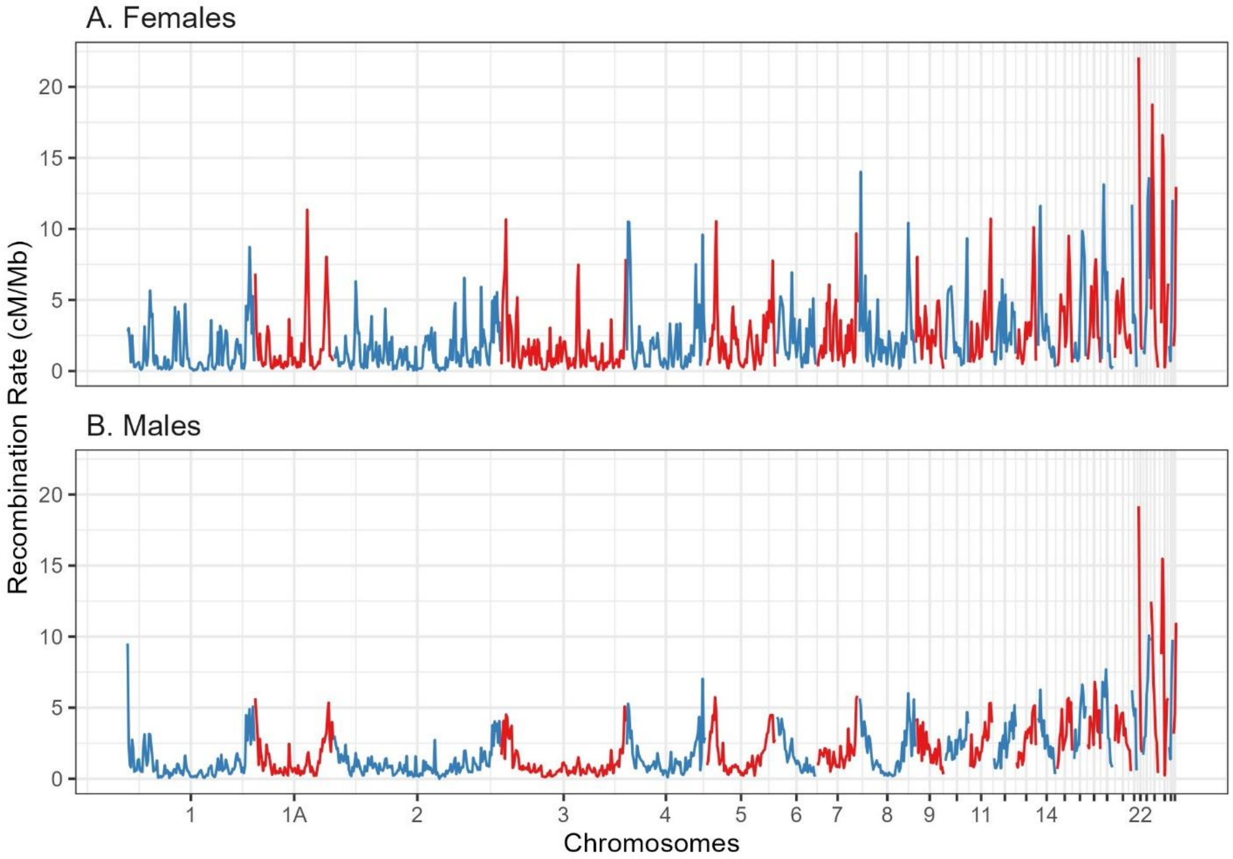


**Figure S2: Recombination rate (cM/Mb) within 1Mb windows in A) female and B) male house sparrows.** Recombination rates per bin were calculated from the linkage map (Table S3) as the difference between the minimum and maximum cM position divided by the difference between the minimum and maximum base pair position. Windows were not included here if they had fewer than 5 SNPs and/or if the difference between the minimum and maximum base pair position was less than 500kb (N = 903 of 930 windows were retained). Lines have been colour-coded by chromosome.

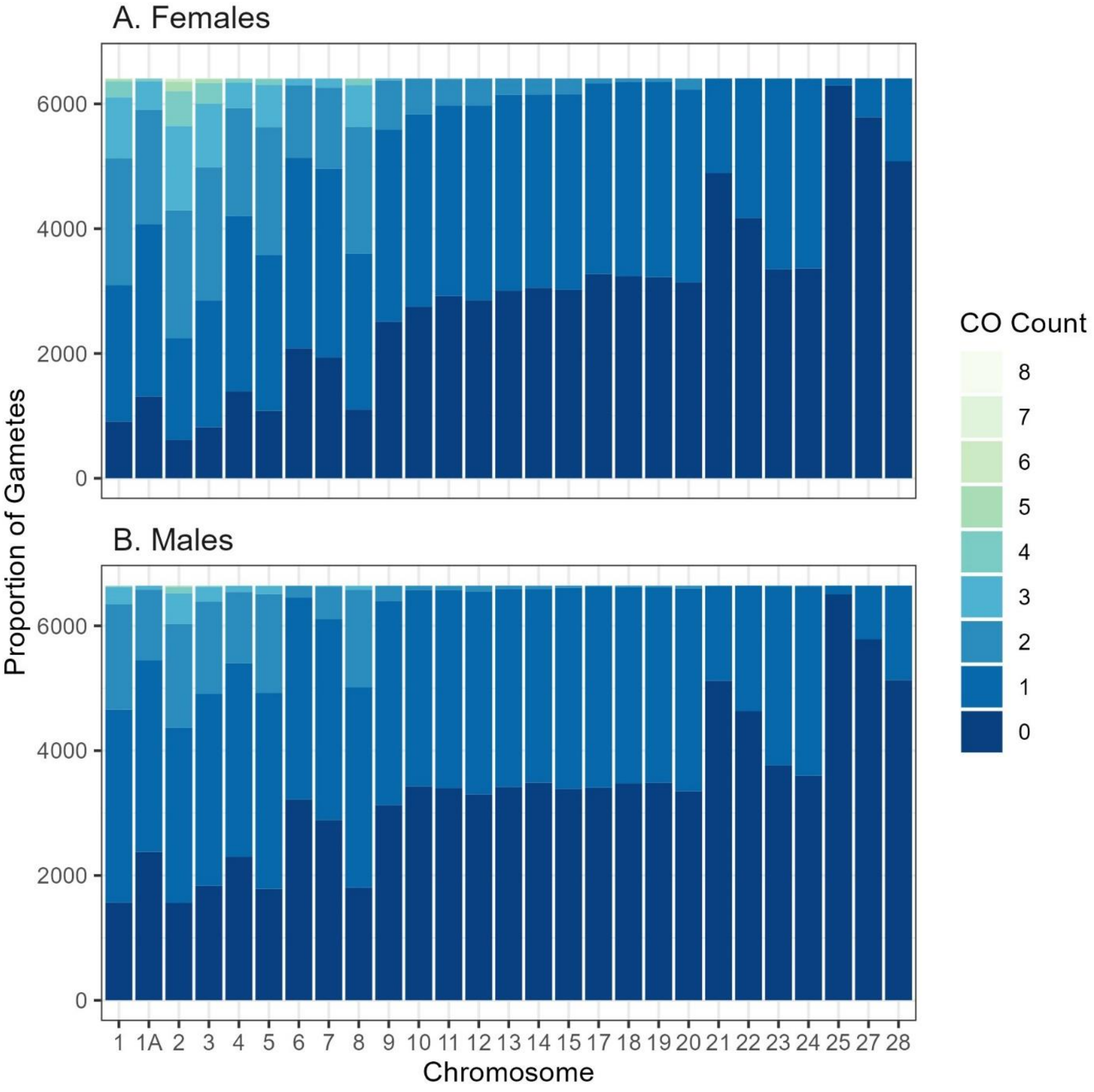


**Figure S3:** Distribution of crossover (CO) counts per chromosome as the proportion of total number of gametes (N = 13,054) faceted by sex. The white dashed line is the minimum expected proportion of gametes with 0 COs per chromosome due to obligate crossing-over and Mendelian segregation of COs into gametes.

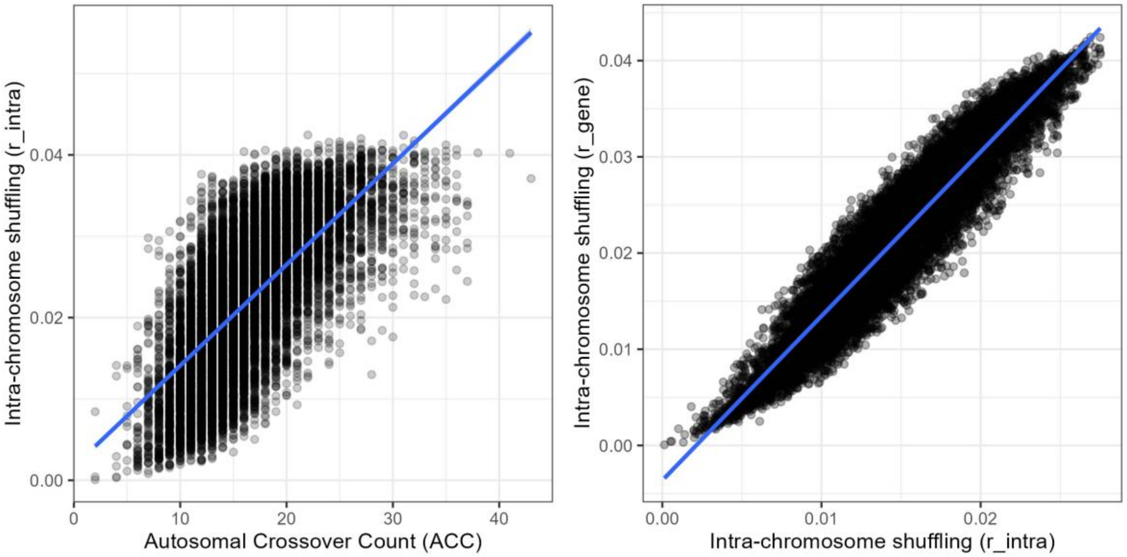


**Figure S4:** Pearson's correlations between A) autosomal crossover count (ACC) and the rate of intra-chromosomal shuffling $\bar{r}_{intra}$ (*$r^2$ = 0.684, P < 0.0001*) and B) between $\bar{r}_{intra}$ and $\bar{r}_{gene}$ ($r^2$ = 0.948, P < 0.001).

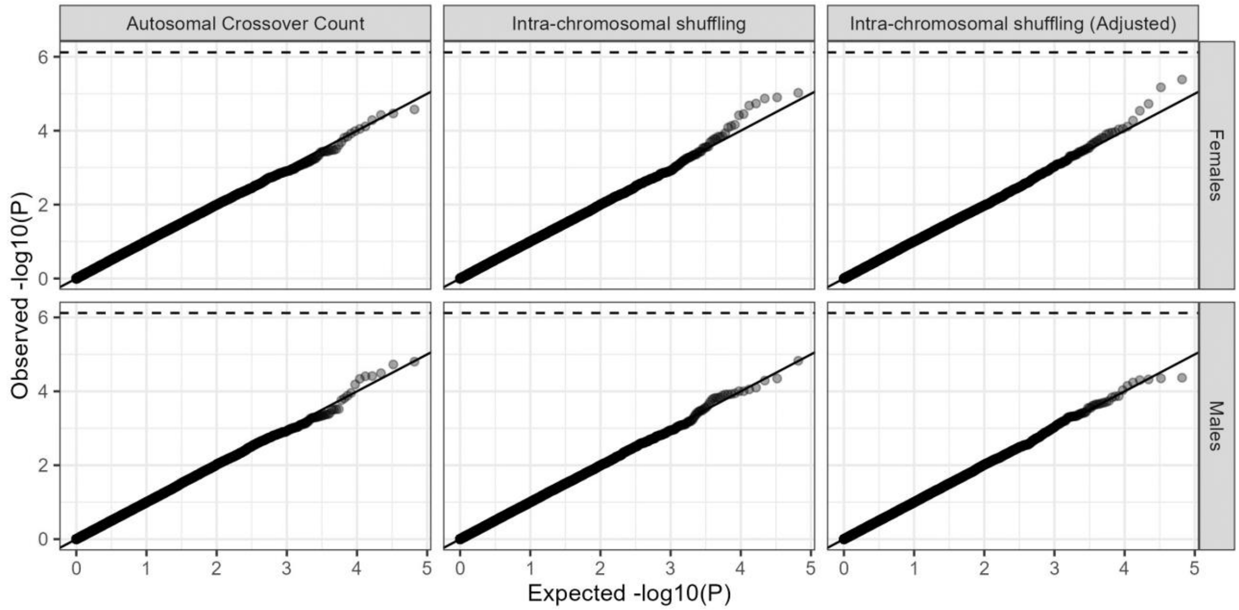


**Figure S5:** P-P plots of the expected (null) distribution of P values for the genome-wide association studies presented in Figure 5 in the main text. The dashed line is the genome-wide significance equivalent to $\alpha = 0.05$. The solid line indicates the Expected = Observed. Association statistics have been corrected with the genomic control parameter $\lambda$.

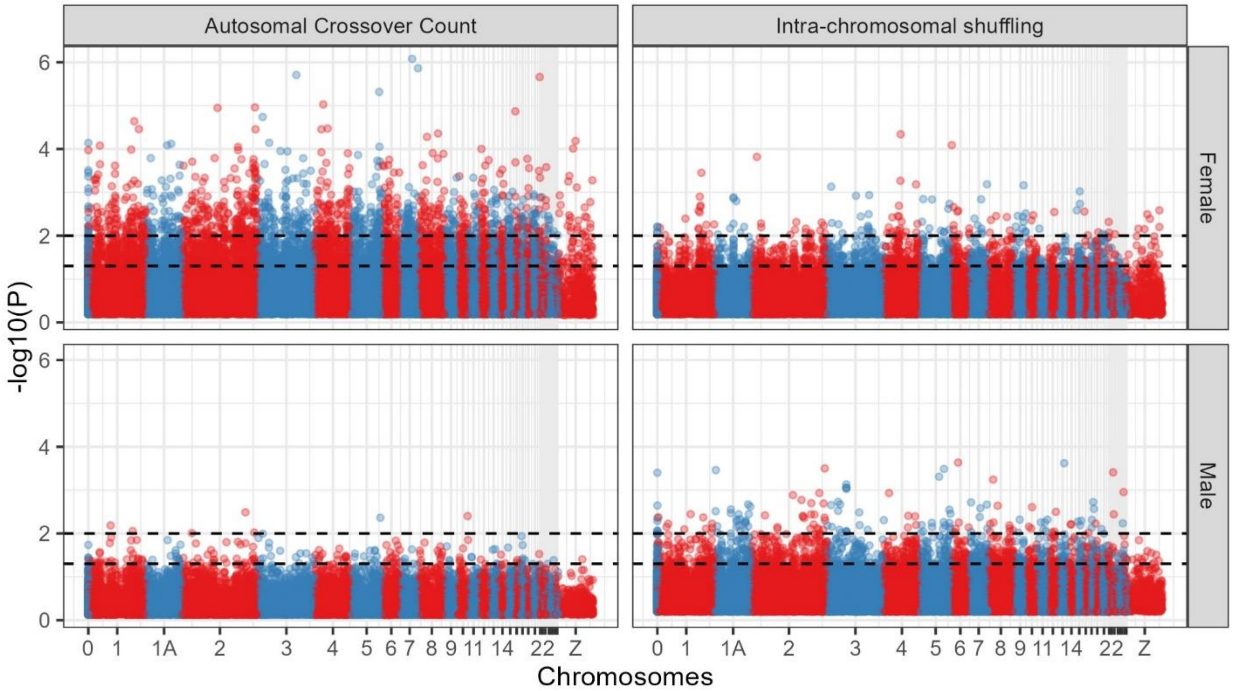


**Figure S6:** Manhattan plots of the significance of the local false sign rate as estimated from ashR Empirical Bayes approach (Stephens 2017). The dashed lines indicate the genome-wide significance equivalent to α = 0.05 (lower) and α = 0.01 (upper). Points have been colour-coded by chromosome.

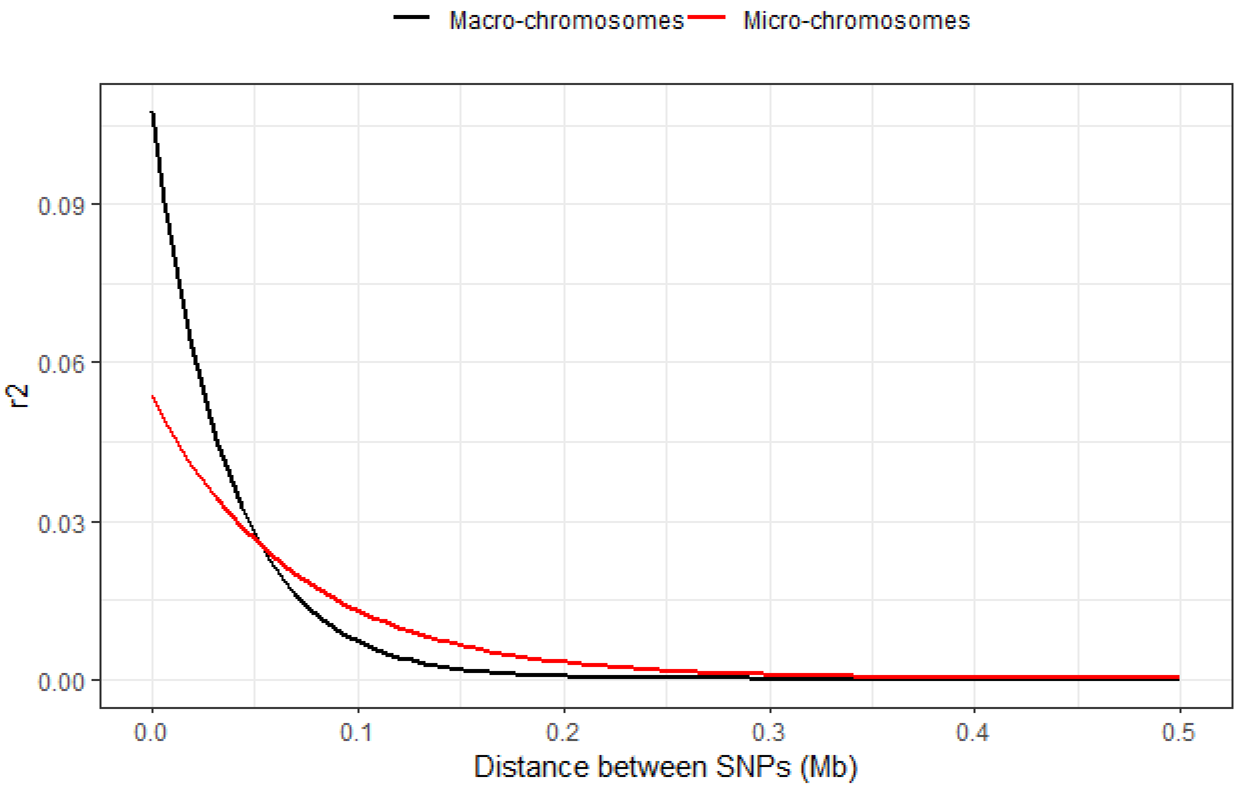


**Figure S7.** Genomic LD decay for SNPs up to 0.5Mb distance. Lines are colour-coded as macro (chromosomes 1A and 1 to 12) and micro (chromosomes 12 to 28). Lines indicate the exponential decay function y ~ a*exp(b *-x) fit in ggplot2 (Wickham 2016).

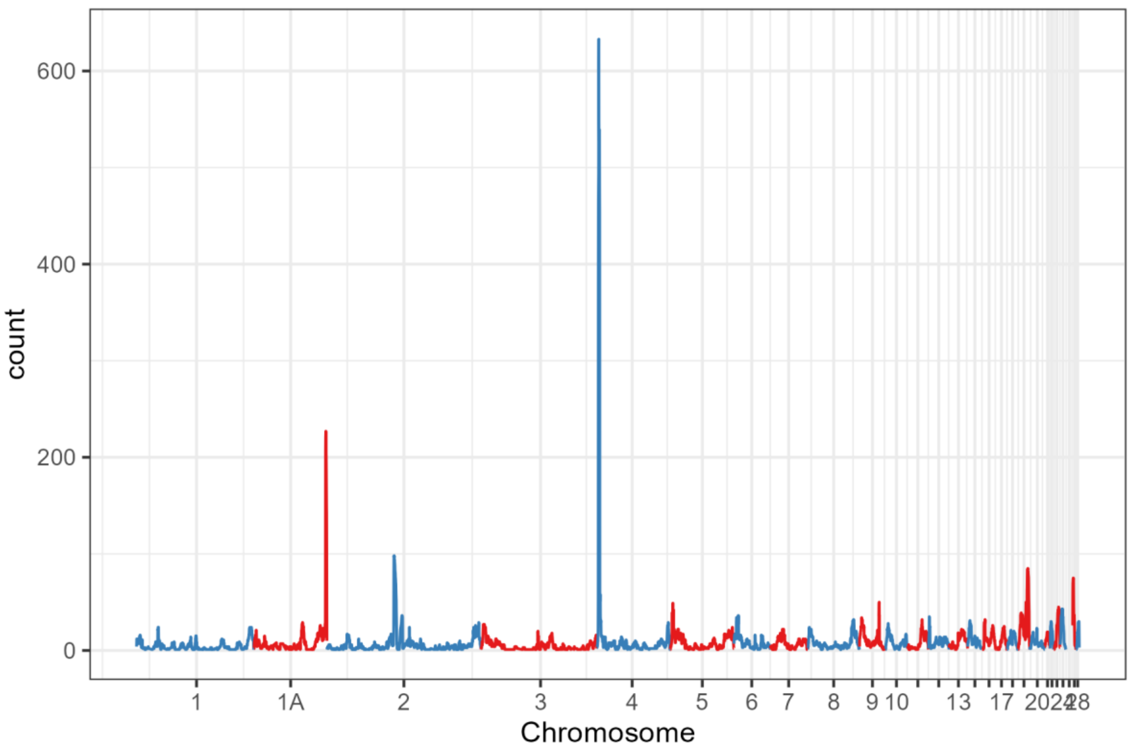


**Figure S8:** Distribution of short double crossover events (<3Mb distance between two crossovers positions) within 100kb genomic intervals along the genome.

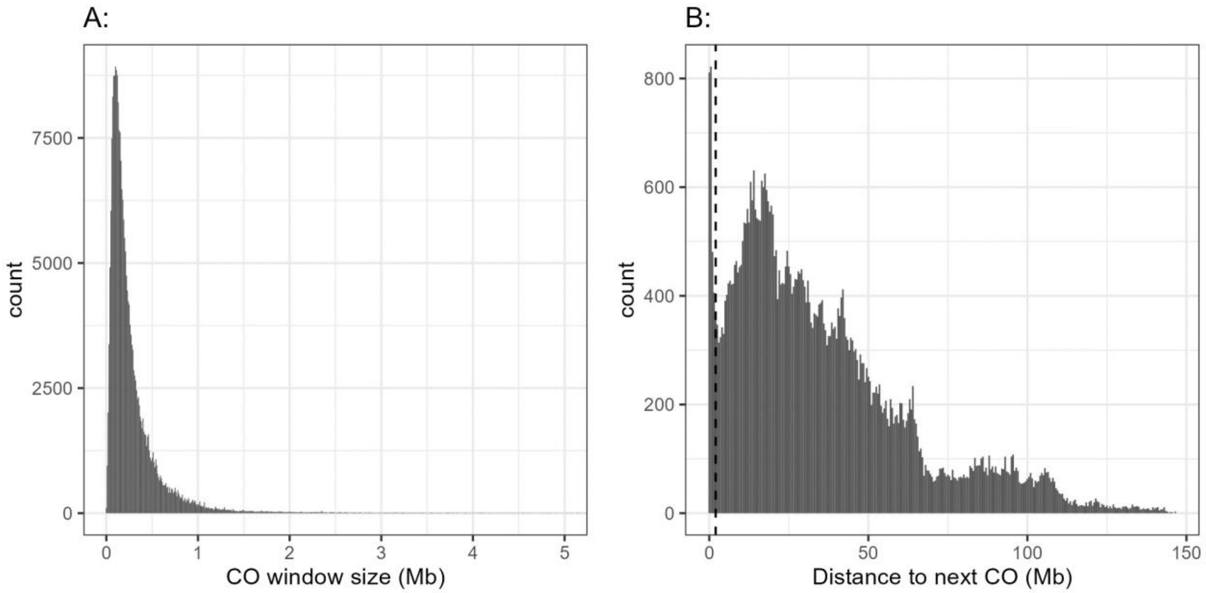


**Figure S9.** The distribution of distances between (A) the start and stop position of crossover (CO) intervals in Mb and (B) the stop position of a given CO and the start position of the next (Mb), in cases where more than one CO occurred on a chromosome in a given meiosis. NB. 2358 data points (0.98% of CO windows) of greater than 5Mb were removed from panel A for ease of visualisation.

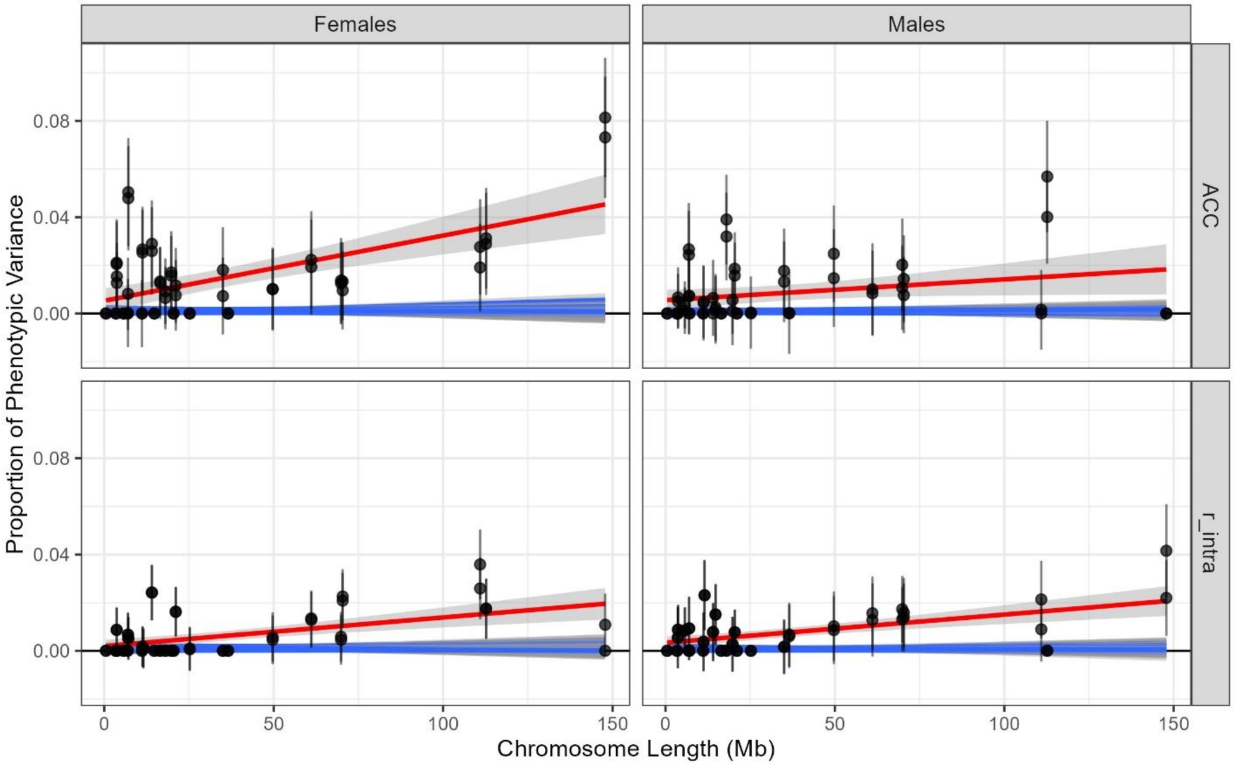


**Figure S10:** Chromosome partitioning of additive genetic effects on autosomal crossover count (ACC) and intra-chromosomal shuffling ($\bar{r}_{intra}$) in females and males. Each point indicates the proportion of phenotypic variance explained by a chromosome-specific genomic relatedness matrix modelled as a function of chromosome length. The red line indicates the true slope from a linear regression, with standard errors. The blue lines indicate the slopes and their standard errors calculated in permuted datasets (N = 100) to test the null distribution of the effects of censoring and heteroscedasticity (Kemppainen and Husby 2018). This analysis showed that this effect is likely to be minimal in the current dataset.

**Table S1:** Measures of autosomal heterochiasmy in birds. This table is adapted from (Malinovskaya *et al.* 2020), who determined the centiMorgan map lengths based on male and female MLH1 foci counts (in cases marked with*). LM = linkage map, CC = crossover count from pedigree data, MLH1 = MLH1 foci and RN = recombination nodules in cytogenetic investigations.

| SPECIES | NAME | FEMALE | MALE | TYPE | DEVIATION | MARKER COUNT | IDS | REFERENCE |
|---|---|---|---|---|---|---|---|---|
| **House sparrow** | *Passer domesticus* | 2240 | 1801 | LM | 0.196 | 6498 | 1898 | (Hagen *et al.* 2020) |
| **House sparrow** | *Passer domesticus* | 1994 | 1627 | LM | 0.184 | 56765 | 2653 | This study |
| **Siberian jay** | *Perisoreus infaustus* | 809 | 694 | LM | 0.142 | 117 | 349 | (Jaari *et al.* 2009) |
| **Zebra finch** | *Taeniopygia guttata* | 899 | 809 | LM | 0.1 | 876 | 354 | (Stapley *et al.* 2008) |
| **Zebra finch** | *Taeniopygia guttata* | 2335 | 2310 | MLH1* | 0.011 | - | - | (Calderón and Pigozzi 2006) |
| **Blue tit** | *Parus caeruleus* | 1046 | 887 | LM | 0.152 | 91 | 525 | (Hansson *et al.* 2010) |
| **Great reed warbler** | *Acrocephalus arundinaceus* | 858 | 552 | LM | 0.357 | 58 | 693 | (Hansson *et al.* 2005) |
| **Collared flycatcher** | *Ficedula albicollis* | 1627 | 1982 | LM | -0.218 | 241 | 322 | (Backström *et al.* 2008) |
| **Collared flycatcher** | *Ficedula albicollis* | 3354 | 3846 | LM | -0.147 | 33627 | 655 | (Kawakami *et al.* 2014) |
| **Great tit** | *Parus major* | 2010 | 1917 | LM | 0.046 | 6554 | 2000 | (van Oers *et al.* 2014) |
| **Honeyeater** | *Lichenostomus melanops* | 1680 | 1864 | LM | -0.110 | 53111 | 257 | (Robledo-Ruiz *et al.* 2022) |
| **Superb fairywren** | *Malurus cyaneus* | 1686 | 1583 | LM | 0.061 | 11414 | 273 | (Peñalba *et al.* 2020) |
| **Barn swallow** | *Hirundo rustica* | 2815 | 2430 | MLH1* | 0.137 | - | - | (Malinovskaya *et al.* 2020) |
| **Domestic chicken** | *Gallus gallus* | 3310 | 3285 | MLH1* | 0.008 | - | - | (Pigozzi 2001) |
| **Domestic chicken** | *Gallus gallus* | 3098 | 3145 | LM | -0.015 | 9268 | ~700 | (Groenen *et al.* 2009) |
| **Domestic chicken** | *Gallus gallus (White Layer)* | 23.9 | 18.2 | CC | 0.313 | 580000 | 1200 | (Weng *et al.* 2019) |
| **Domestic chicken** | *Gallus gallus (Brown Layer)* | 31.3 | 29.4 | CC | 0.065 | 42000 | 5108 | (Weng *et al.* 2019) |
| **Domestic goose** | *Anser anser* | 3655 | 3030 | MLH1* | 0.171 | - | - | (Torgasheva and Borodin 2017) |
| **Japanese quail** | *Coturnix japonica* | 2815 | 2815 | MLH1* | 0 | - | - | (Calderón and Pigozzi 2006) |
| **Pale martin** | *Riparia diluta* | 2380 | 2450 | MLH1* | -0.029 | - | - | (Malinovskaya *et al.* 2020) |
| **Pigeon** | *Columba livia* | 3135 | 3235 | RN* | -0.032 | - | - | (Pigozzi and Solari 1999) |
| **Turkey** | *Meleagris gallopavo* | 2077 | 2431 | LM | -0.17 | 531 | 1043 | (Aslam *et al.* 2010) |

**Table S2:** Summary data for the house sparrow with sex-averaged and sex-specific linkage maps. Full linkage map data is provided in Table S2.

| Chromosome | Female length (cM) | Male length (cM) | Chromosome length (Mb) | Number of SNP loci |
|---|---|---|---|---|
| **1** | 154.068 | 118.812 | 112673038 | 6289 |
| **1A** | 117.594 | 94.704 | 69872796 | 4336 |
| **2** | 199.294 | 145.428 | 147816218 | 8774 |
| **3** | 162.744 | 111.689 | 110917403 | 6510 |
| **4** | 137.781 | 102.373 | 70339410 | 4713 |
| **5** | 130.924 | 102.813 | 61080839 | 4243 |
| **6** | 81.6 | 56.161 | 35005501 | 2515 |
| **7** | 86.547 | 64.304 | 36520066 | 2734 |
| **8** | 123.032 | 91.404 | 49673226 | 3135 |
| **9** | 68.033 | 53.064 | 25208338 | 1766 |
| **10** | 59.8 | 48.081 | 21074797 | 1450 |
| **11** | 57.894 | 50.799 | 20430685 | 1546 |
| **12** | 55.632 | 48.834 | 19767051 | 1392 |
| **13** | 51.731 | 47.14 | 18005134 | 1152 |
| **14** | 49.291 | 43.147 | 16462598 | 1007 |
| **15** | 49.995 | 44.806 | 14030935 | 899 |
| **17** | 42.472 | 42.385 | 11239481 | 593 |
| **18** | 43.807 | 44.033 | 11524157 | 736 |
| **19** | 45.259 | 44.373 | 11088665 | 661 |
| **20** | 48.183 | 46.624 | 14763639 | 1034 |
| **21** | 22.722 | 21.305 | 5690223 | 343 |
| **22** | 32.993 | 28.014 | 3636148 | 102 |
| **23** | 40.842 | 39.029 | 6974644 | 197 |
| **24** | 38.971 | 38.756 | 7069745 | 198 |
| **25** | 14.302 | 12.837 | 476344 | 9 |
| **26** | 43.488 | 47.547 | 6890404 | 175 |
| **27** | 14.026 | 16.521 | 3681783 | 108 |
| **28** | 21.385 | 22.233 | 3406084 | 148 |
| **Total** | 1994.41 | 1627.216 | 915319352 | 56765 |

**Table S3:** High density linkage map for House Sparrows estimated using Lep-MAP v3. Mb is the chromosome position in bp relative to the Passer_domesticus-1.0 genome assembly (Elgvin *et al.* 2017). Female_cM and Male_cM indicate the female and male linkage map positions in cM, respectively, using the Morgan mapping function.
[File: TableS3_House_Sparrow_Linkage_Map.txt]

**Table S4:** Fixed effect results for each recombination rate measure from animal models. Wald statistics are analogous to a $\chi^2$ test with 1 degree of freedom. Recombination measures are taken for chromosomes 1, 1A to 20. Random effects are reported in Table 1 in the main text.

| Measure | Sex | Fixed Effect | Solution | Standard error | Z Ratio | Wald statistic | P ($\chi^2$) |
|---|---|---|---|---|---|---|---|
| **ACC** | Female | Intercept | 128.454 | 75.7 | 1.697 | 40875.003 | 0 |
| | | Total_Coverage | -3.09E-07 | 1.76E-07 | -1.759 | 70.374 | 0 |
| | | $Total\_Coverage^2$ | 2.09E-16 | 1.02E-16 | 2.046 | 4.184 | 0.0408 |
| | Male | Intercept | 47.622 | 66.046 | 0.721 | 53227.815 | 0 |
| | | Total_Coverage | -1.11E-07 | 1.53E-07 | -0.73 | 55.62 | 8.79E-14 |
| | | $Total\_Coverage^2$ | 8.22E-17 | 8.81E-17 | 0.933 | 0.87 | 0.351 |
| **r_intra** | Female | Intercept | 0.114 | 0.116 | 0.987 | 50423.708 | 0 |
| | | Total_Coverage | -2.53E-10 | 2.69E-10 | -0.941 | 28.3 | 1.04E-07 |
| | | $Total\_Coverage^2$ | 1.73E-19 | 1.56E-19 | 1.112 | 1.236 | 0.266 |
| | Male | Intercept | -0.195 | 0.152 | -1.278 | 15956.602 | 0 |
| | | Total_Coverage | 4.78E-10 | 3.52E-10 | 1.358 | 1.455 | 0.228 |
| | | $Total\_Coverage^2$ | -2.69E-19 | 2.03E-19 | -1.326 | 1.757 | 0.185 |
| **r_intra** | Female | Intercept | -0.006 | 0.092 | -0.066 | 95699.678 | 0 |
| **(Corrected for ACC)** | | ACC | 0.001 | 1.61E-05 | 54.686 | 2990.536 | 0 |
| | | Total_Coverage | 3.83E-11 | 2.15E-10 | 0.179 | 54.443 | 1.6E-13 |
| | | $Total\_Coverage^2$ | -2.26E-20 | 1.24E-19 | -0.182 | 2.356 | 0.125 |
| | Male | Intercept | -0.234 | 0.127 | -1.84 | 26512.466 | 0 |
| | | ACC | 0.001 | 2.40E-05 | 51.215 | 2623.013 | 0 |
| | | Total_Coverage | 5.69E-10 | 2.94E-10 | 1.936 | 2.762 | 0.0966 |
| | | $Total\_Coverage^2$ | -3.43E-19 | 1.7E-19 | -2.023 | 1.971 | 0.16 |

**Table S5:** GWAS and Empirical Bayes false discovery results for all loci with non-zero effects on autosomal crossover count (ACC) and intra-chromosomal shuffling ($\bar{r}_{intra}$). MAF is the minor allele frequency. *effB* is the slope of allelic effects and *se_effB* is the standard error of the slope, as estimated using RepeatABEL. *chi2.1df* is the association statistic and *Pc1df* is the corrected P-value after genomic control. *NegativeProb* and *PositiveProb* are the probabilities of negative and positive sign of the slope. *lfsr* and *lfdr* are the local false sign rate and local false discovery rates, respectively. The *qvalue* is the significance of the effect. *PosteriorMean* and *PosteriorSD* are the posterior mean and standard deviations, respectively.

**[File: TableS5_Sig_GWAS_and_FDR_Results.txt]**

**Table S6.** Linear regressions and correlations of chromosome partitioning of additive genetic effects on autosomal crossover count (ACC) and intra-chromosomal shuffling ($\bar{r}_{intra}$) in females and males. Each model regresses the proportion of phenotypic variance explained by a chromosome-specific genomic relatedness matrix modelled as a function of chromosome length. The variance explained is modelled for recombination rates estimated on all chromosomes (1-20 & 1A), which accounts for *cis* and *trans* effects combined, or for recombination rates excluding that chromosome, accounting for *trans* effects only. The underlying raw data is provided in Table S7.

| Measure | Sex | Model | Effect | Estimate | SE | t | P | Adj R2 |
|---|---|---|---|---|---|---|---|---|
| ACC | Females | cis & trans | Intercept | 5.65E-03 | 3.62E-03 | 1.561 | 0.131 | 0.328 |
| | | | Length | 2.65E-04 | 7.17E-05 | 3.703 | **0.0011** | |
| | | trans | Intercept | 4.89E-03 | 3.81E-03 | 1.284 | 0.211 | 0.323 |
| | | | Length | 2.76E-04 | 7.54E-05 | 3.664 | **0.0012** | |
| | Males | cis & trans | Intercept | 4.90E-03 | 3.32E-03 | 1.475 | 0.153 | 0.067 |
| | | | Length | 1.12E-04 | 6.58E-05 | 1.697 | 0.102 | |
| | | trans | Intercept | 6.12E-03 | 2.93E-03 | 2.087 | 0.047 | 0.004 |
| | | | Length | 6.12E-05 | 5.81E-05 | 1.053 | 0.302 | |
| r_intra | Females | cis & trans | Intercept | 1.47E-03 | 1.90E-03 | 0.772 | 0.447 | 0.325 |
| | | | Length | 1.44E-04 | 3.84E-05 | 3.743 | **9.12E-04** | |
| | | trans | Intercept | 2.43E-03 | 1.92E-03 | 1.266 | 0.217 | 0.155 |
| | | | Length | 9.46E-05 | 3.87E-05 | 2.44 | **0.022** | |
| | Males | cis & trans | Intercept | 2.59E-03 | 1.90E-03 | 1.367 | 0.183 | 0.382 |
| | | | Length | 1.61E-04 | 3.82E-05 | 4.203 | **2.75E-04** | |
| | | trans | Intercept | 4.11E-03 | 1.60E-03 | 2.568 | 0.016 | 0.133 |
| | | | Length | 7.32E-05 | 3.22E-05 | 2.271 | **0.032** | |

**Table S7.** Full chromosome partitioning results of additive genetic effects on autosomal crossover count (ACC) and intra-chromosomal shuffling ($\bar{r}_{intra}$) in females and males for each chromosome, considering models of all recombination measures (cis & trans) or recombination on all other chromosomes. The estimate and SE (standard error) are the proportion of phenotypic variance explained by the effect. Models included the additive genetic effect (for all chromosomes excluding the chromosome), the additive genetic effect of that chromosome, the permanent environment effect (repeated measures) and the residual variance.

**[File: TableS7_Chr_h2_FULL_results.txt]**

### Referenced Literature


Aslam M. L., J. W. M. Bastiaansen, R. P. M. A. Crooijmans, A. Vereijken, H.-J. Megens, *et al.*, 2010 A SNP based linkage map of the turkey genome reveals multiple intrachromosomal rearrangements between the turkey and chicken genomes. BMC Genomics 11: 647.

Backström N., N. Karaiskou, E. H. Leder, L. Gustafsson, C. R. Primmer, *et al.*, 2008 A gene-based genetic linkage map of the collared flycatcher (Ficedula albicollis) reveals extensive synteny and gene-order conservation during 100 million years of avian evolution. Genetics 179: 1479–1495.

Calderón P. L., and M. I. Pigozzi, 2006 MLH1-focus mapping in birds shows equal recombination between sexes and diversity of crossover patterns. Chromosome Res. 14: 605–612.

Elgvin T. O., C. N. Trier, O. K. Tørresen, I. J. Hagen, S. Lien, *et al.*, 2017 The genomic mosaicism of hybrid speciation. Sci. Adv. 3: e1602996.

Groenen M. A. M., P. Wahlberg, M. Foglio, H. H. Cheng, H.-J. Megens, *et al.*, 2009 A high-density SNP-based linkage map of the chicken genome reveals sequence features correlated with recombination rate. Genome Res. 19: 510–519.

Hagen I. J., S. Lien, A. M. Billing, T. O. Elgvin, C. Trier, *et al.*, 2020 A genome-wide linkage map for the house sparrow (Passer domesticus) provides insights into the evolutionary history of the avian genome. Mol. Ecol. Resour. 20: 544–559.

Hansson B., M. Akesson, J. Slate, and J. M. Pemberton, 2005 Linkage mapping reveals sex-dimorphic map distances in a passerine bird. Proc. Biol. Sci. 272: 2289–2298.

Hansson B., M. Ljungqvist, D. A. Dawson, J. C. Mueller, J. Olano-Marin, *et al.*, 2010 Avian genome evolution: insights from a linkage map of the blue tit (Cyanistes caeruleus). Heredity (Edinb.) 104: 67–78.

Jaari S., M.-H. Li, and J. Merilä, 2009 A first-generation microsatellite-based genetic linkage map of the Siberian jay (Perisoreus infaustus): insights into avian genome evolution. BMC Genomics 10: 1.

Kawakami T., L. Smeds, N. Backström, A. Husby, A. Qvarnström, *et al.*, 2014 A high-density linkage map enables a second-generation collared flycatcher genome assembly and reveals the patterns of avian recombination rate variation and chromosomal evolution. Mol. Ecol. 23: 4035–4058.

Kemppainen P., and A. Husby, 2018 Inference of genetic architecture from chromosome partitioning analyses is sensitive to genome variation, sample size, heritability and effect size distribution. Mol. Ecol. Resour. 18: 767–777.

Malinovskaya L. P., K. Tishakova, E. P. Shnaider, P. M. Borodin, and A. A. Torgasheva, 2020 Heterochiasmy and sexual dimorphism: The case of the barn swallow (Hirundo rustica, hirundinidae, Aves). Genes (Basel) 11: 1119.

Oers K. van, A. W. Santure, I. De Cauwer, N. E. M. van Bers, R. P. M. A. Crooijmans, *et al.*, 2014 Replicated high-density genetic maps of two great tit populations reveal fine-scale genomic departures from sex-equal recombination rates. Heredity (Edinb.) 112: 307–316.

Peñalba J. V., Y. Deng, Q. Fang, L. Joseph, C. Moritz, *et al.*, 2020 Genome of an iconic Australian bird: High-quality assembly and linkage map of the superb fairy-wren (Malurus cyaneus). Mol. Ecol. Resour. 20: 560–578.

Pigozzi M. I., and A. J. Solari, 1999 Equal frequencies of recombination nodules in both sexes of the pigeon suggest a basic difference with eutherian mammals. Genome 42: 315–321.

Pigozzi M. I., 2001 Distribution of MLH1 foci on the synaptonemal complexes of chicken oocytes. Cytogenet. Cell Genet. 95: 129–133.

Robledo-Ruiz D. A., H. M. Gan, P. Kaur, O. Dudchenko, D. Weisz, *et al.*, 2022 Chromosome-length genome assembly and linkage map of a critically endangered Australian bird: the helmeted honeyeater. Gigascience 11. https://doi.org/10.1093/gigascience/giac025

Stapley J., T. R. Birkhead, T. Burke, and J. Slate, 2008 A linkage map of the zebra finch Taeniopygia guttata provides new insights into avian genome evolution. Genetics 179: 651–667.

Stephens M., 2017 False discovery rates: a new deal. Biostatistics 18: 275–294.

Torgasheva A. A., and P. M. Borodin, 2017 Immunocytological Analysis of Meiotic Recombination in the Gray Goose (Anser anser). Cytogenet. Genome Res. 151: 27–35.

Weng Z., A. Wolc, H. Su, R. L. Fernando, J. C. M. Dekkers, *et al.*, 2019 Identification of recombination hotspots and quantitative trait loci for recombination rate in layer chickens. J. Anim. Sci. Biotechnol. 10: 20.

Wickham H., 2016 About the ggplot2 Package. J. Appl. Comput. Math. 5. https://doi.org/10.4172/2168-9679.1000321
